# Concussive head trauma perturbs axon initial segment function in axotomized and intact layer 5 pyramidal neurons

**DOI:** 10.1101/2022.04.29.490079

**Authors:** Alan C. Harris, Jianli Sun, Kimberle Jacobs

## Abstract

The axon initial segment (AIS) is a critical locus of control of action potential (AP) generation and neuronal information synthesis. Concussive traumatic brain injury gives rise to diffuse axotomy, and the majority of neocortical axonal injury arises at the AIS. Consequently, concussive traumatic brain injury might profoundly disrupt the functional specialization of this region. To investigate this hypothesis, one and two days after mild central fluid percussion injury in Thy1-YFP-H mice, we recorded high-resolution APs from axotomized and adjacent intact layer 5 pyramidal neurons and applied a second derivative (2°) analysis to measure the AIS- and soma-regional contributions to the AP upstroke. All layer 5 pyramidal neurons recorded from sham animals manifested two stark 2° peaks separated by a negative intervening slope. In contrast, within injured mice, we discovered a subset of axotomized layer 5 pyramidal neurons in which the AIS-regional 2° peak was abolished, a functional perturbation associated with diminished excitability, axonal sprouting and distention of the AIS as assessed by staining for ankyrin-G. Our analysis revealed an additional subpopulation of both axotomized and intact layer 5 pyramidal neurons that manifested a melding together of the AIS- and soma-regional 2° peaks, suggesting a more subtle aberration of sodium channel function and/or translocation of the AIS initiation zone closer to the soma. When these experiments were repeated in animals in which cyclophilin-D was knocked out, these effects were ameliorated, suggesting that trauma-induced AIS functional perturbation is associated with mitochondrial calcium dysregulation.

## Introduction

During concussive, or mild, traumatic brain injury (mTBI), acceleration of the brain within the cranial vault sets into motion a complex sequence of cellular pathogenesis, comprising dysregulated neuronal ionic gradients, decreased glycolysis, formation of free radical species, and accumulation of calcium in mitochondria (Blennow *et al*., 2012; Prins *et al*., 2013). Furthermore, influx of calcium ions into the axoplasm stimulates calpain-mediated proteolysis of neurofilaments and voltage-gated sodium channels (McGinn *et al*., 2009; von Reyn *et al*., 2009; Brocard *et al*., 2016). Neurofilament proteolysis in conjunction with stretch-mediated disassembly of microtubules blocks the flow of anterograde protein trafficking and gives rise to areas of organelle accumulation, termed retraction bulbs (Povlishock & Christman, 1995). These bulbs swell, and ultimately the distal boundary of the bulb disconnects from the axon 6-12 hours after initial injury (Greer *et al*., 2013). This process of traumatic axotomy is pronounced among layer 5 cortical pyramidal neurons, which constitute the principal source of electrochemical outflow from the neocortex.

The axon initial segment (AIS) is an unmyelinated subdomain of the neuron just distal to the axon hillock. Owing to low regional capacitance (Kole & Stuart, 2012), a high density of voltage-gated sodium channels (Kole *et al*., 2008), and to the presence of the Na_v_1.6 isoform (Hu *et al*., 2009), which has a relatively hyperpolarized voltage of activation, the distal AIS is a specialized locus in which the action potential (AP) has the lowest threshold to initiation. Because the AIS controls AP generation, the tuning of its conductance properties and morphology represents a potent mechanism toward the maintenance of neuronal homeostasis (Evans *et al*., 2013). Plasticity of the length of the AIS, the position of the AIS relative to the soma, and the intrinsic membrane properties of the AIS have been observed in response to sensory deprivation and enrichment (Kuba *et al*., 2010; Kuba *et al*., 2015; Jamann *et al*., 2021), aging and embryonic development (Gutzmann *et al*., 2014; Atapour & Rosa, 2017), and neurological and psychiatric disease (Hinman *et al*., 2013; Marin *et al*., 2016; Yermakov *et al*., 2018). It has been demonstrated that the AIS is especially vulnerable to axotomy secondary to mTBI, with the majority of axotomy arising at the AIS and para-AIS (Greer *et al*., 2013); furthermore, various models of TBI give rise to contraction of the distal end of the AIS of intact neurons as measured by staining for ankyrin-G (Baalman *et al*., 2013; Vascak *et al*., 2017). This distal contraction is associated with a reduction of AIS-regional voltage acceleration two days after TBI (Vascak *et al*., 2017). Given the critical specialization of the AIS toward the control of AP generation and neuronal information synthesis, the phenomenon of diffuse axonal injury might disrupt information outflow from the neocortex and consequently perturb sensory perception and global cognitive function.

Following generation at the AIS, the AP conducts in the orthodromic direction in a saltatory manner to the nodes of Ranvier. Simultaneously, the AP back-propagates to the somatodendritic region, where it may modulate dendritic conductances (Stuart *et al*., 1997). There exists a period of latency between these events, which manifests as a transient lag between two peaks upon analysis of the second derivative (2°) of the voltage with respect to time during the AP upstroke. It has been demonstrated by means of regional blockade of sodium channels in hippocampal pyramidal cells (Meeks & Mennerick, 2007) and cerebellar Purkinje cells (Khaliq & Raman, 2006) that the first of these peaks corresponds to the depolarization of the AIS, and the second peak corresponds to the depolarization of the somatodendritic region. In the present study, we replicate this result in layer 5 pyramidal neurons and use this analysis as the basis of an assay of the functional integrity of the AIS and soma. We demonstrate that all layer 5 pyramidal neurons within the sham condition manifest both peaks. On the other hand, after mTBI, the first of these peaks is abolished in a subset of axotomized layer 5 pyramidal neurons 1- and 2-days post-injury, a functional perturbation associated with a significantly reduced frequency-current slope and diminished staining for ankyrin-G. Additionally, we describe a subset of intact layer 5 pyramidal neurons within the vicinity of axotomy that manifest an AIS-regional peak that melds into the soma-regional peak. This waveform may represent a trauma-induced aberration of sodium channel function that exists even within morphologically undamaged neurons.

## Methods

### Experimental Animals

Two mouse lines were used to generate the data of the present study. The first line expresses Thy1-driven yellow fluorescent protein (YFP) within a subset of layer 5 pyramidal neurons (Feng *et al*., 2000) and was acquired from Jackson Labs [B6Cg-TgN(Thy1-YFPH) 2Jrs, stock number 003782; Bar Harbor, ME, USA]. Ear punches were inspected by fluorescence microscopy to confirm the presence of the YFP transgene. To generate the second line, these same YFP-H mice were crossed with Cyclophilin-D (CypD) knock-out mice, which have been shown to be more resistant to the assembly and opening of the mitochondrial permeability transition pore and the consequent derangement of intracellular calcium dynamics and mitochondrial swelling (Basso *et al*., 2005; Forte *et al*., 2007). The gene encoding CypD is ppif, and thus these knock-out mice are also known as ppif^- /-^. Here the YFP-H strain is referred to as wildtype (WT), and the CypD knock-out mice crossed with the YFP-H are referred to as CypDKO. For this study, male mice (P56-P90) were used. The mice were group housed in 12 h/12 h non-reversed light cycle conditions on corn cob bedding with access to food and water *ad libitum*. All animal procedures were approved by the institutional animal care and use committee of Virginia Commonwealth University.

### Central fluid percussion injury (cFPI)

Mild cFPI was induced as described previously (Greer *et al*., 2012; Hånell *et al*., 2015; Rowe *et al*., 2016; Sun & Jacobs, 2016). Mice were anesthetized with 4% isoflurane in 100% O_2_, with anesthesia maintained with 2% isoflurane during surgery. The body temperature was thermostatically controlled via a heating pad (Harvard Apparatus, Holliston, MA, USA) and maintained at 37°C. A pulse oximetry sensor (STARR Life Sciences, Oakmont, PA, USA) was used to monitor pulse rate, respiratory rate, and blood oxygenation intraoperatively. A 3.0 mm circular craniectomy was made along the sagittal suture midway between bregma and lambda. Care was taken to avoid any insult to the underlying dura. The cFPI injury at this location consistently produces diffuse axonal injury throughout the primary somatosensory cortex. A sterile Leur-Loc syringe hub cut from a 20-gauge needle was affixed to the craniectomy site using cyanoacrylate and dental cement then filled with sterile saline. This procedure required 45-75 minutes. The animal was then removed from anesthesia and monitored in a warmed cage until fully ambulatory (60–80 min of recovery). Injury or sham procedure was applied thereafter.

For injury induction, each animal was re-anesthetized with 4% isoflurane in 100% O_2_ for four minutes, and the hub was filled with sterile saline and attached to a fluid percussion apparatus (Custom Design and Fabrication; Virginia Commonwealth University; Richmond, VA). A mild severity injury (1.7 ± 0.06 atmospheres) was induced by a brief fluid pressure pulse upon the intact dura. The peak pressure was measured by the transducer (Tektronix 5111) and displayed on an oscilloscope. After injury, the animals were monitored visually for loss of spontaneous respiration or the emergence of clonus. Transient derangement of respiratory rhythm that resolved into eupnea was routinely observed. The duration of transient unconsciousness was measured by the time interval between the injury and the return of the spontaneous righting reflex. While sham righting times occur within around 1 minute, within this cFPI injury paradigm, injury is considered effectively mild when righting occurs within eight minutes, a standard that held true for all injured mice included here. After recovery of the righting reflex, animals were placed in a warmed holding cage and monitored during recovery (typically ∼60 min) before being returned to the vivarium. For sham injury, all the above steps were followed except for the release of the pendulum to induce the injury.

### Acute slice preparation and patch-clamp recording

One or two days after injury or sham, mice were deeply anesthetized with isoflurane and perfused through the aorta with semi-frozen, sucrose substituted artificial cerebrospinal fluid (aCSF) previously saturated by a mixture of 95% O_2_ / 5% CO_2_ (carbogen) containing (in mM): 2.5 KCl, 10 MgSO_4_, 0.5 CaCl_2_, 1.25 NaH_2_PO_4_, 234 sucrose, 11 glucose, and 26 NaHCO_3_. Subsequently the mice were decapitated, and the brain was rapidly extracted and chilled in this same aCSF. Scrupulous care was undertaken to generate anatomically coronal slices in which vibratome-induced axotomy was minimized. To this end, the brain was carefully docked within an acrylic mouse brain slicer matrix (Zivic Instruments) to generate anatomically coronal slices. 300-*μ*m thick coronal slices were cut with a Leica VT 1200 vibratome (Leica Microsystems, Wetzlar, Germany). These slices were incubated for 30 min at 34°C in aCSF containing (in mM): 126 NaCl, 3 KCl, 2 MgCl_2_, 2 CaCl_2_, 1.25 NaH_2_PO_4_, 10 glucose, and 26 NaHCO_3_, bubbled continuously with carbogen. The slices remained at room temperature thereafter until transferred to the recording chamber, which was maintained at 30±0.5° C. The slices were secured by a harp, and a 60X water-immersion objective was used to visually identify YFP^+^ layer 5 pyramidal neurons of primary somatosensory cortex ventral to the injury site. Neurons possessing an axon that could be traced along its trajectory into the subcortical white matter were considered intact, while those terminating in an axonal bleb were considered axotomized. Care was taken to select only neurons axotomized deep to the surface of the slice in order to exclude any neurons axotomized by the vibratome during slicing. Among YFP+ layer 5 pyramidal neurons in the YFP-H condition, differentiating between these two morphologies is readily feasible within the living brain slice (Greer *et al*., 2012).

Within the recording chamber, slices were perfused with aCSF bubbled continuously with carbogen. Patch electrodes (final resistance 2–4 MΩ) were pulled from borosilicate glass (World Precision Instruments, Sarasota, FL, USA). The intracellular solution contained (in mM): 130 K-gluconate, 10 Hepes, 11 EGTA, 2.0 MgCl2, 2.0 CaCl2, 4 Na-ATP, 0.2 Na-GTP, 0.4% biocytin. Electrode capacitance was electronically compensated. Recordings were made via a MultiClamp 700B (Molecular Devices, Sunnyvale, CA, USA) and digitized via a Digidata 1440A and pClamp software (Molecular Devices).

During current clamp recordings, the membrane potential was maintained at -60 mV with continuous depolarizing or hyperpolarizing current as needed. Access resistance was continuously monitored. If the access resistance increased by 15% at any time between break in and the culmination of data collection, an attempt was made to clear the pipette tip via gentle positive pressure. If this attempt was not successful, the recording was terminated. Most neurons had a resting membrane potential (RMP) between -60 mV and -70 mV (Table 1), except for a few severely axotomized neurons that rested between -60 mV and -55 mV. The axotomized neurons for both 1- and 2-day survival times were significantly more depolarized than those of the 2-day sham condition (Table 1). On the measure of input resistance, there was no significant difference in subject groups or experimental conditions (WT or CypDKO) and no significant interaction between these two (Table 1).

**Table 1.** Resting membrane potential (RMP) and input resistance (Rin) for all groups. Subject groups are listed in the first column while experimental condition was wildtype or CypDKO. # = significant, with a Bonferroni post-hoc for RMP showing that Sham 2D was significantly different from TBI AX1D (p = 0.025) and TBI AX2D (p = 0.022). In addition, for the RMP measure, TBI IN2D was significantly different from TBI AX1D (p = 0.048) and TBI AX2D (p = 0.039).

| Subject Group | RMP (mV) |  | Rin (MΩ) |  |
| --- | --- | --- | --- | --- |
|  | WT | CypDKO | WT | CypDKO |
| **Sham 1D** | -66.6 ±1.1 (22) | -66.3 ±1.2 (7) | 78.7 ±8.8 (22) | 57.4 ±3.4 (7) |
| **Sham 2D** | -68.9 ±0.8 (17) | -65.6 ±1.9 (7) | 71.3 ±5.4 (17) | 61.1 ±6.3 (9) |
| **TBI AX1D** | -62.8 ±0.7 (20) | -67.2 ±1.7 (9) | 73.0 ±9.5 (21) | 74.8 ±6.1 (9) |
| **TBI AX2D** | -62.9 ±1.3 (13) | -64.7 ±2.6 (6) | 66.1 ±6.8 (15) | 61.0 ±7.2 (6) |
| **TBI IN1D** | -64.4 ±0.6 (22) | -65.3 ±1.6 (11) | 77.5 ±10.4 (23) | 63.9 ±3.4 (11) |
| **TBI IN2D** | -68.0 ±1.9 (14) | -67.4 ±1.4 (8) | 78.0 ±6.6 (14) | 62.4 ±4.3 (8) |
| ***p-values*** |  |  |  |  |
| ***Subject group*** | 0.048<sup>#</sup> |  | 0.970 |  |
| ***Experimental Condition*** | 0.551 |  | 0.080 |  |
| ***Interaction*** | 0.113 |  | 0.785 |  |

Two main protocols of current clamp recordings were run. The first protocol consisted of a brief injection of depolarizing current, titrated to evoke 1-2 APs, which were recorded unfiltered and digitized at 200 kHz (high resolution APs for 2° analysis). This procedure was repeated 5-9 times at 0.2 Hz. To identify intrinsic membrane properties and the pattern and kinetics of AP discharge, the second protocol consisted of a series of 400 msec long hyperpolarizing and depolarizing current injections, which were low pass filtered with a 10kHz cut-off and digitized at 20 kHz.

In order to correlate the biphasic components of the AP upstroke to the regional depolarization of the AIS and the soma, a patch pipette containing tetrodotoxin (TTX, 2 µM) was positioned approximately 30 µm distal to the axon hillock (to dissociate the AIS component) or over the soma (to dissociate the somatic component) of visualized YFP^+^ layer 5 pyramidal neurons within the WT sham condition. Subsequently, TTX was applied through the patch pipette via a Picospritzer (Parker Hannifin, Mayfield Heights, Ohio, USA) at 30 psi for a duration of 8 msec and at a rate of 0.1 Hz. Immediately after each puff of TTX, an AP was evoked via square wave depolarizing current injection and subject to 2° analysis. The slice was positioned within the chamber with the apical dendrites projecting toward the aCSF inflow and the axon projecting toward the outflow to minimize TTX spillage onto the soma during TTX application to the AIS.

### Post hoc immunohistochemistry

After each recording, neurons were depolarized slowly to – 40 mV, and the patch pipette was gradually withdrawn over the course of approximately twenty seconds to re-seal the neuronal membrane. Slices were fixed in 4% paraformaldehyde for 70 minutes then washed six times in 0.1M PBS and stored in 0.1M PBS at 4° C until processing. To reconstruct the morphology of recorded neurons, slices were washed three times for 5 minutes with 0.1M PBS then reacted overnight at 4° C with a solution of 0.1 M PBS containing Texas Red-conjugated streptavidin (1:500) in the presence of 0.5% triton-X. To visualize the structure of the AIS, slices were then treated with heat-based epitope retrieval. Slices were immersed within a solution of 10 mM sodium citrate dihydrate (294.10 g/mol; pH = 8.0) and heated on a block to 80° C for 10 minutes. The slices were allowed to cool for 10 minutes, then blocked in a solution of 10% normal horse serum and 0.5% triton-X within 0.1M PBS at room temperature for 1 hour. Slices were then washed three times with 0.1M PBS for 10 minutes and incubated in a solution containing anti-ankyrin-G primary antibody (1:400; mouse IgG2a, clone N106/36, NeuroMab) for 24 hours at 4°C. The slices then were washed three times with 0.1M PBS and reacted with a solution containing the secondary antibody (Alexa Fluor 405; 1:500) for 2 hours at room temperature. Subsequently, slices were washed three times for 10 minutes in 0.1M PBS and mounted on glass slides with Vectashield non-hardening anti-fade mounting medium, and the slides sealed with clear nail polish. Slices thus prepared were imaged at 63X via a Zeiss 880 laser scanning confocal microscope. Serial scanning of each of the three channels (Texas Red, YFP, and Alexa Fluor 405) was employed, and care was taken to image the entire axon of all successfully recovered neurons. For measurement of the distance to axotomy, the distance from the emergence of the axon to the site of the first retraction bulb was measured using the freehand line tool on the biocytin or YFP channel within ImageJ. For axotomized neurons of club-shaped morphology and lacking classical retraction bulbs, the distance from the emergence of the axon to the end of the axon was measured. For measurement of the AIS regional axonal width, the width of the axon of neurons axotomized more than 30µm from the soma was measured at 30µm using the straight-line tool within ImageJ. For neurons axotomized within 30µm of the soma, the width was measured at its widest point along the axon excluding the retraction bulb itself.

### Data analysis

Data analysis was performed with Clampfit software (Molecular Devices, Sunnyvale, CA, USA) and custom programs in Visual Basic for Applications, within Microsoft Excel (Microsoft, Redmond, WA, USA). For the APs generated for 2° analyses, in all cases only the first AP in the sweep was analyzed. The digitized AP was filtered with a 7-point boxcar filter repeated three times, prior to calculation of the first derivative and the 2°. AP threshold was measured as the membrane potential at 10 mV/msec. For each cell typically 5 APs were analyzed (3-5APs), with measurements made of each individual AP and then those measurements averaged to create the record per cell.

Unless otherwise specified, significance was tested using one or two-way ANOVAs, with Bonferroni post hoc test (SPSS, IBM). Results are reported as mean ±SEM.

## Results

To probe for the existence of trauma-induced perturbation of AIS function, we recorded from ex-vivo brain slices prepared from mice subjected to mild cFPI 1 or 2 days prior to experimentation. In the resulting slices, we used a 60X water immersion objective to follow the trajectory of the axons of YFP^+^, layer 5 pyramidal neurons to entrance into the subcortical white matter (intact neurons), or to the axonal bleb/swelling (axotomized neurons, Greer *et al*., 2012). Neurons thus visualized were targeted for whole cell patch clamp recording. From these neurons, high-resolution APs were generated by a brief depolarizing current injection at a level that typically evoked 1-2 APs. In addition, the membrane voltage response to a series of hyperpolarizing and depolarizing pulses of 400 msec duration in order to identify the AP firing pattern and other intrinsic cellular properties was recorded. We note that all YFP^+^, layer 5 pyramidal neurons tested possessed some degree of the hyperpolarization-activated cyclic nucleotide-gated current (I_h_), which manifests as a depolarizing sag in response to hyperpolarizing current injection. The amplitude of this sag was added to the amplitude of the depolarizing afterpotential in the manner described by Gee and colleagues (2012). It was determined that all neurons recorded in this study could be classified as Type A (Supplemental Fig. 1).

The 2° analysis of the AP upstroke has been demonstrated to contain two components (Cp1, Cp2), which have been shown to represent the sequential depolarization of the AIS and the soma, respectively, in CA3 pyramidal neurons (Meeks & Mennerick, 2007) and cerebellar Purkinje cells (Khaliq & Raman, 2006). To extend this analysis to layer 5 neocortical pyramidal neurons and to confirm that our methods allowed the differentiation of AP generation at the AIS and the soma, APs were recorded before, during and after the focal application of TTX (2 µM) to either the axon at the expected location of the AIS or to the soma (Fig. 1A, B). In sham controls, the 2° of the AP always contained two components with clearly separate peaks (Fig. 1 C). Application of TTX to the axon approximately 30 *μ*m distal to the axon hillock reduced the peak amplitude of Cp1 to a greater degree than that of Cp2 (Fig. 1C-E). Continued application of TTX produced APs with a gradually diminishing Cp1, which ultimately was abolished outright, giving rise to a monophasic waveform. Within seconds of cessation of TTX application, 2° analysis again demonstrated two peaks. This process of eliminating the first peak then allowing its recovery could be repeated within the same neuron. Prior to elimination of Cp1, TTX application also decreased the time between component peaks and depolarized the AP threshold potential (Fig. 1 E, F). Just prior to elimination of Cp1, the separation of the peaks became less apparent, such that Cp1 and Cp2 began to meld into one another, giving rise to a shoulder and a peak (Fig. 1C, 20 sec). To quantify this transition from two peaks to a single peak, we used a new measure in which the 2° was first normalized to the second (or only) peak. A line was then drawn from 99% to 30% of the second (or only) peak (green lines in Fig. 1C). The difference between the actual data and this theoretical line was then calculated for each point and summed for all points creating the summed difference from linear slope (LS_diff_). In cases of 2 clear peaks separated by a negative intervening slope, the summed difference from this linear slope was always greater than 2.25. The LS_diff_ dropped below this threshold in cases of a shoulder+peak and in cases of a single peak. These results were consistent in all neurons tested, whether from sham controls or the IN1D group (Fig. 1G-K).

**Figure 1.**
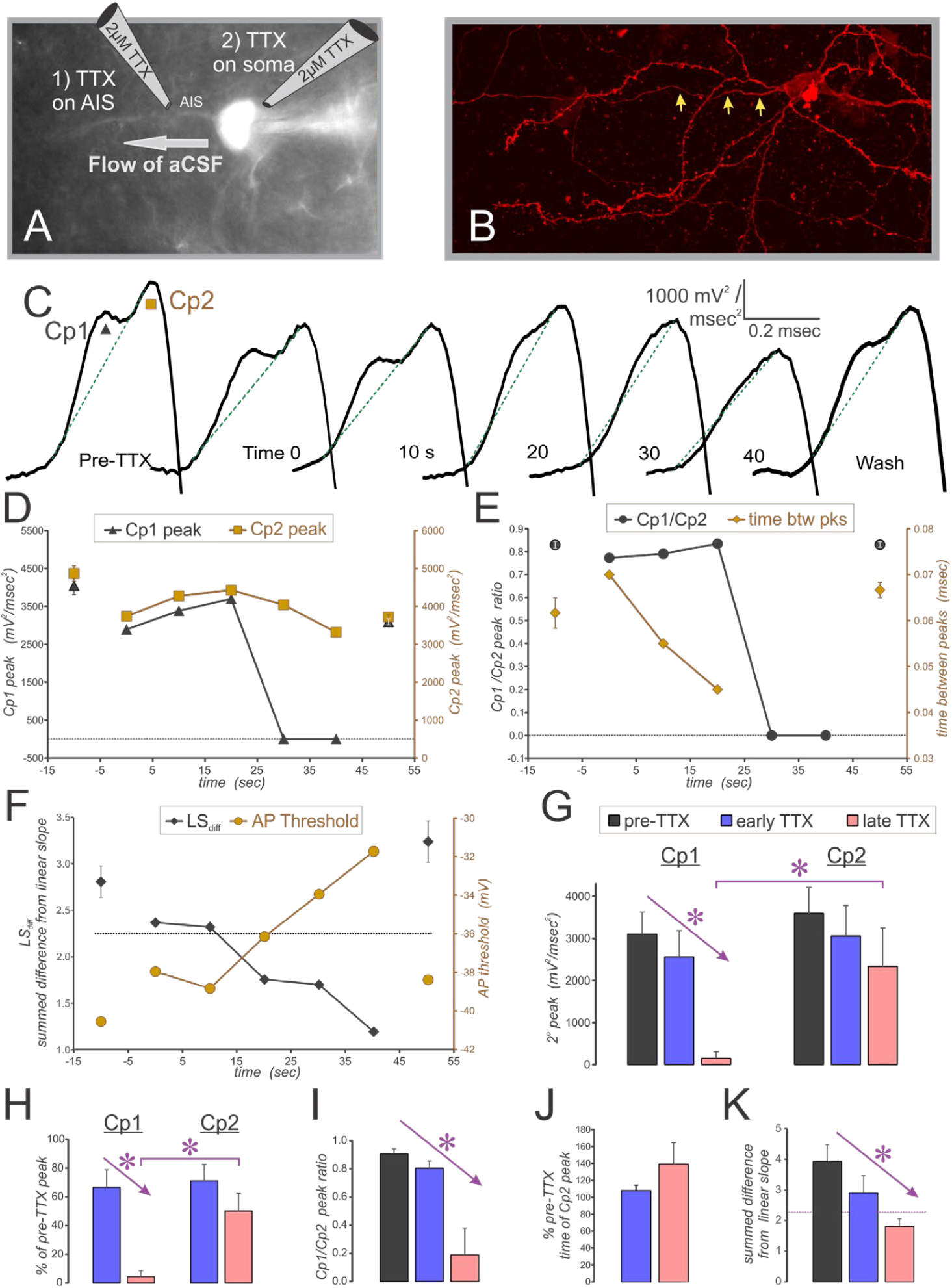
The biphasic components of the AP upstroke, visualized with a 2° analysis are caused by sequential depolarization of the AIS and the soma. Results shown here are for application of TTX to the AIS. **A**. Experimental schematic, including image taken during recording. TTX is puffed onto the AIS (position 1) or the soma (position 2, see Fig. 2). **B**. Maximum projection of the confocal image for same cell shown in A. Arrows indicate axon. **C**. 2° of the AP from the cell shown in A & B, during sequential application of TTX. Green dashed lines show linear slope from 99% to 10% of Cp2 peak. Symbols in first panel show the associated peaks for D. **D-F**. Measurements made on the same cell as A-C. The pre-TTX and wash measurements are each an average of 5, while individual measurements of TTX application are shown. Black markers are plotted on the left axis and brown on the right axis. Shown in F is the summed difference points from the real data to a line drawn from the 99% of the second peak to 10% of the peak. Note the dashed line for the left axis that indicates a threshold level of 2.25 dividing 2° with 2 peaks vs a shoulder + peak or single peak. **G**. Decrement of Cp1 with TTX application was typical of control cells (N = 3), with some decrement of the Cp2 peak but significantly more for the Cp1 peak. A 2-way repeated measures ANOVA showed a significant effect of TTX timing and of the interaction of TTX location (AIS vs soma) with the TTX timing. **H-K**. A similar effect was found in 2 intact cells one day after injury that had 2 peaks pre-TTX. Here all 5 cells are grouped together. In H the peaks are normalized to the pre-TTX level. This shows the relatively selective effect on the Cp1 peak of the TTX applied to the AIS. Here also a repeated measures 2-way ANOVA showed a significant effect of TTX timing and of the interaction between TTX location (with results in figure 2 considered) and TTX timing. **I**. The ratio of Cp1/Cp2 peaks also shows a significant reduction, with repeated measures 1-way ANOVA. **J**. The time of the Cp2 peak was not significantly different after TTX. **K**. The LSdiff measure falls below the 2.25 threshold, indicating a single peak, at the late TTX time point, and this measure is significantly different with TTX application (repeated measures ANOVA).

Movement of the TTX puffer to the location of the soma reduced the amplitude of Cp2, but typically did not eliminate it (Fig. 2). Because the puffer electrode tip size was small relative to the soma, the entire soma could not be covered by TTX application; thus, we did not expect to abolish Cp2 outright. While targeting the soma, some spillage of TTX onto the region of the AIS did occur, as the aCSF flowed toward the AIS. Consequently, Cp1 was also affected; however, the effect was greatest on Cp2 (Fig. 2C, D). This result can be appreciated upon normalization of the 2° to the Cp1 peak (Fig. 2B). TTX application to the soma gave rise to the converse effects compared to TTX application to the AIS on measures of the Cp1/Cp2 ratio, time between peaks and LS_diff_. Furthermore, there evolved a deepening of the valley between the Cp1 and Cp2 peaks, consistent with an increase in the spatial separation between each zone of activation. To quantify this phenomenon, we measured the minimum value between peaks after normalization to the Cp1 peak (Fig. 2E, right axis, brown squares, 2G). The reduction of this minimum was typical for all neurons tested with TTX application to the soma. From these results, we considered the 2° Cp1 to represent depolarization of the AIS and Cp2 to represent depolarization of the soma. Thus, hereafter Cp1 will be referred to as the AIS-regional peak acceleration of the membrane potential, and Cp2 will be referred to as the soma-regional peak acceleration of the membrane potential.

**Figure 2.**
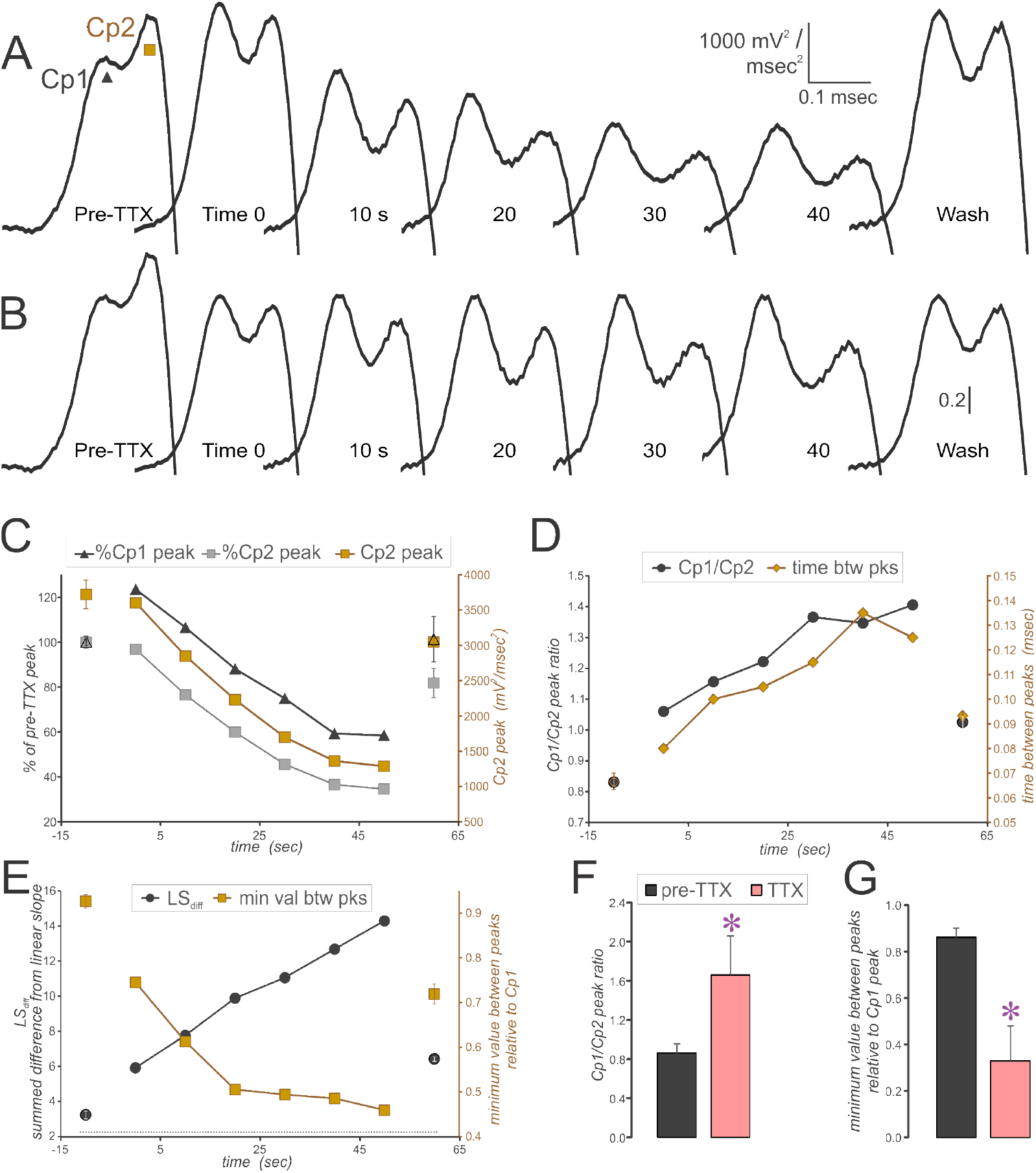
Effect of TTX applied to the soma of a layer 5 pyramidal neuron. **A**. 2° of the AP recorded before, during and after TTX application to the soma. **B**. Same waveforms as shown in A after normalization to the Cp1 peak, showing the relative effect on Cp2. Data in C-E are from this cell, with averages of 5 evoked APs measured for pre-TTX and Wash points. **C**. Left axis shows percent of the pre-TTX peak for Cp1 (black) and Cp2 (gray). Right axis shows Cp2 peak (brown). **D**. Because the TTX applied to the soma has a greater effect on Cp2 than Cp1, the Cp1/Cp2 ratio increases with each TTX application (left axis, black). Time between Cp1 and Cp2 peaks also increases (right axis, brown). **E**. The LSdiff (black, left axis) increases, while the minimum value between peaks relative to Cp1 peak (brown, right axis) decreases with TTX application to the soma. **F**. The effect on the Cp1/Cp2 ratio showed a consistent increase (t-test, 1 tailed, p<0.05, n = 3). **G**. The minimum value between peaks relative to Cp1 peak (t-test, p<0.05, n = 3).

Injection of square wave depolarizing current gave rise to a qualitatively similar AP in sham controls and in axotomized and intact neurons 1 and 2 days after TBI (Fig. 3). Calculation of the first derivative of the AP revealed a change in slope of the AP rise to peak for sham neurons compared to some of those in the TBI groups (Fig. 3B). This result was more evident when plotted against the membrane potential and when compared to the 2° (Fig. 3C, D). When the 2° contained two peaks, a notch could typically be seen in the plot of the first derivative versus membrane potential, indicating a distinct threshold potential for AIS versus soma activation zones. Because concussive TBI-induced axotomy manifests at the AIS within the majority of axotomized neurons (Greer *et al*., 2013), we expected that the AIS function would be perturbed in axotomized neurons, especially those axotomized at the region of the AIS. We discovered a subset of the AX1D and AX2D neurons in which the AIS-regional peak was abolished, although other neurons of these groups manifested 2 peaks similar to those within shams (see examples in Figs. 3, 4). Morphologically, most of the neurons with a single peak were characterized by a wider, distended AIS and diminished staining intensity of ankyrin-G compared to neighboring neurons (Fig. 4, 1D, 2D, also see 4G). Within the AX1D and AX2D groups—which contained neurons with 2 peaks, with shoulder+peak and with a single peak—measurements were made on the biocytin-filled neurons (Fig. 4F-H). While most of the 1-peaked neurons were axotomized in the vicinity of the soma (Fig. 4F), some of the 2-peaked neurons were axotomized within the AIS and 40 *μ*m or less from the soma, suggesting that intracellular signaling cascades rather than mechanical disruption *per se* affect that ability of the AIS to generate an AP after trauma. The width of the AIS was measured 30*μ*m from the soma for neurons axotomized beyond 30*μ*m, while for those axotomized within 30*μ*m of the soma, the width was measured at its widest point along the axon excluding the retraction bulb (Fig. 4G). The 1-peaked 2° neurons had a significantly wider AIS compared to the 2-peaked and shoulder+peak neurons (1-way ANOVA, Bonferroni post-hoc, p<0.05).

**Figure 3.**
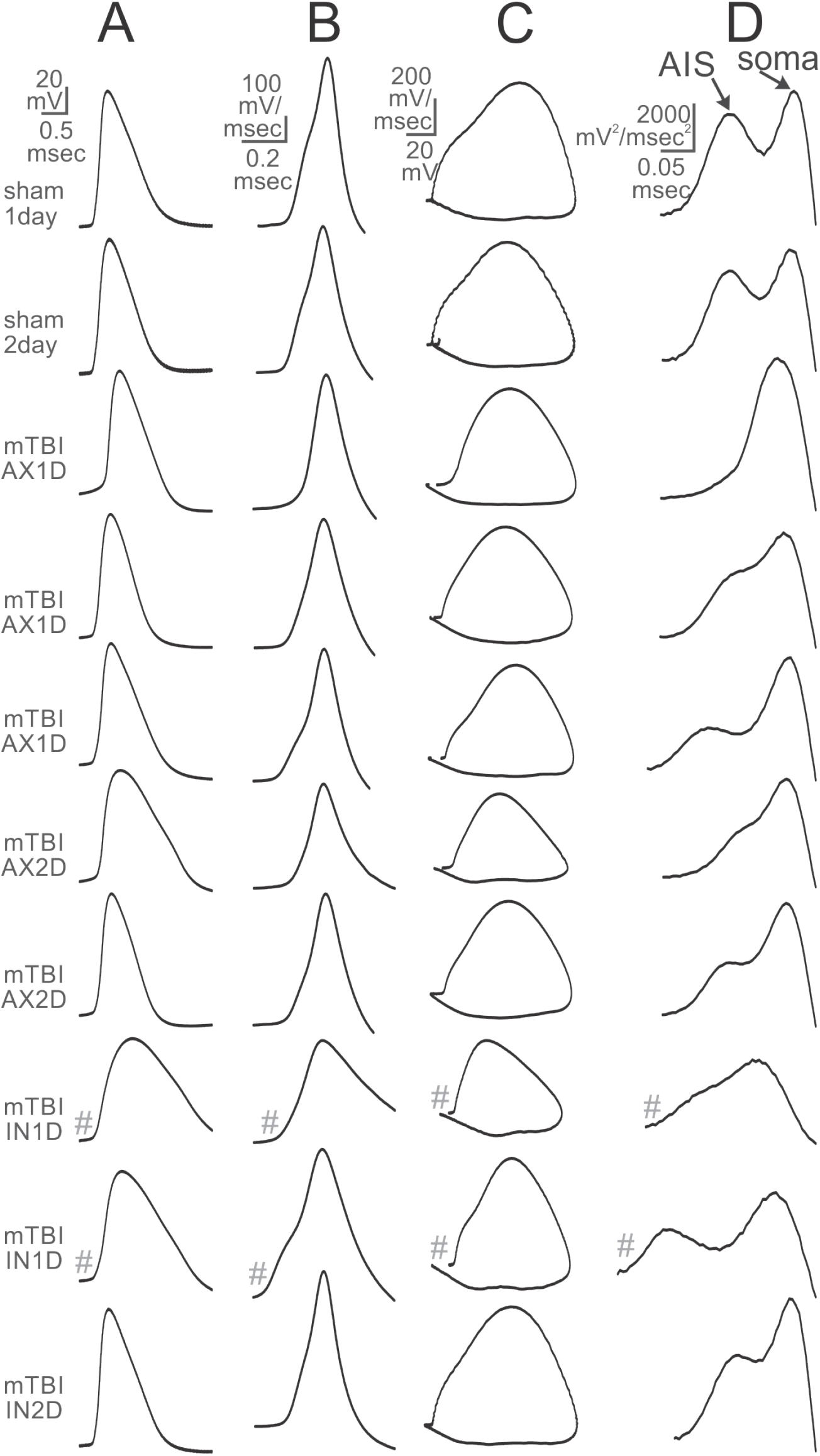
Examples of recordings from each subject group. Waveforms labeled with # indicate that they are shown at 2X vertical scale. A. AP produced by brief depolarizing pulse. B. First derivative of the AP shown in A. C. First derivative as in B but plotted against membrane potential. D. 2° of AP shown in A. All shams had two peaked 2°, some neurons from the mTBI AX1 and 2D groups had single peaked 2°, while others had a shoulder and then a peak, and some were 2 peaked. The mTBI IN1 and 2D groups had some cells with a shoulder and then a peak and some clear 2 peaked 2°.

**Figure 4.**
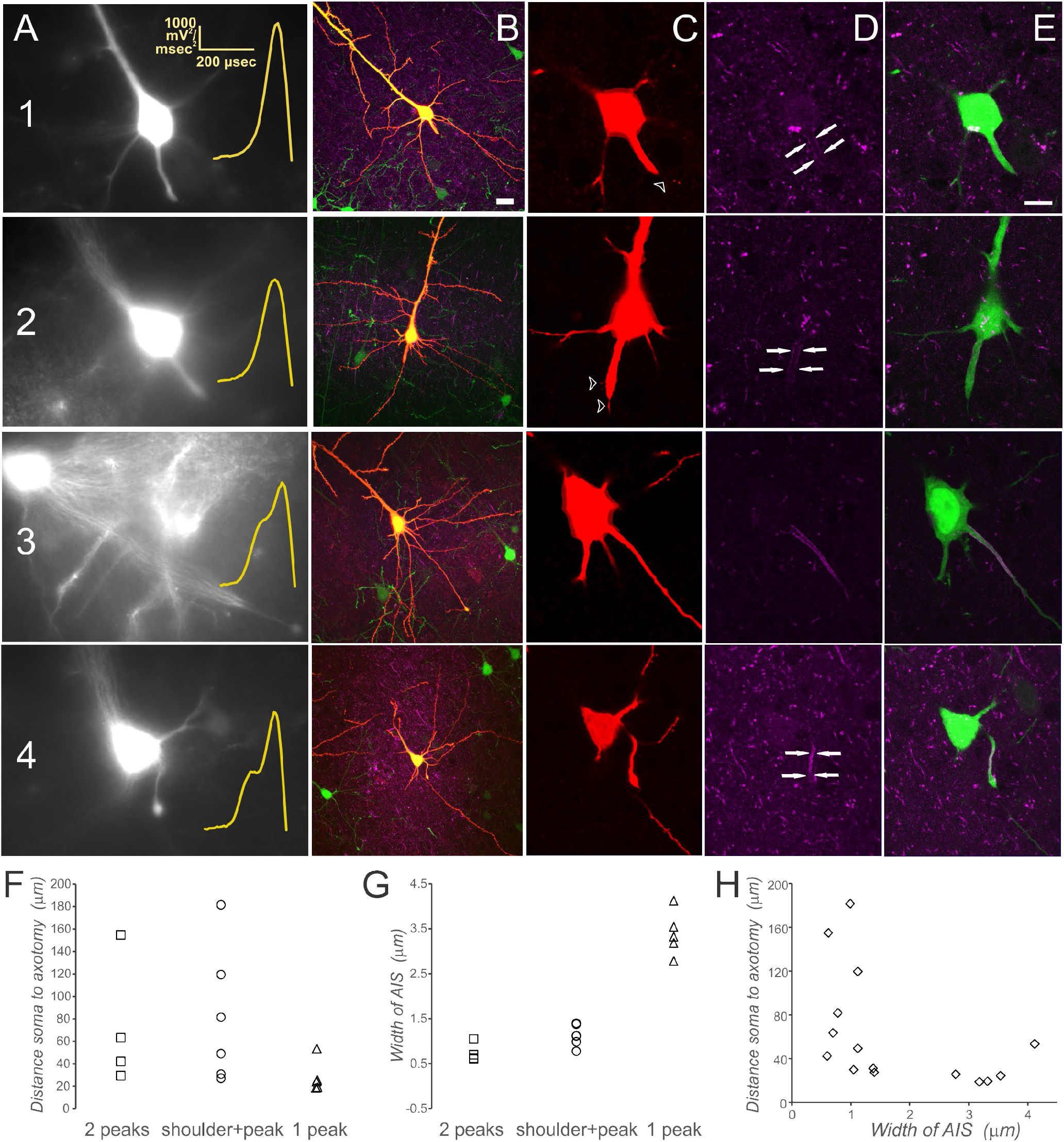
AIS morphology and distance to axotomy. Examples of three neurons axotomized near the soma (1, 2, and 4) and one far from the soma (3). **A**. Targeted neurons as visualized by endogenous YFP expression within the living brain slice just prior to recording. The inset in yellow shows the 2° of the AP recorded from the same cell. **B**. Maximum intensity projections of the same neurons after recovery of biocytin (red) and immunohistochemical staining of ankyrin-G (violet). Endogenous YFP fluorescence is green. **C-E**. Morphological details of the axon initial segments of neurons 1-4 imaged at 63X. Arrows within D indicate the axon initial segments of recorded neurons as visualized by immunohistochemical labeling of ankyrin-G, while E depicts the merged image of ankyrin-G and endogenous YFP. Note that neurons 1 and 2 manifest a diminished intensity of staining for ankyrin-G and a wider, distended AIS. Note also the filipodia-like sprouts emanating from the AIS of neurons 1 and 2 (arrowheads in C). These morphological properties correlate with an absence of discernable AIS-regional voltage acceleration during the AP upstroke. Neuron shown in 1 is the same as that from which recordings are shown in Fig. 3C, while neuron 3 is the same as Fig. 3F, and neuron 4 is the same as Fig. 3D. Scalebar in 1B = 20 µm, for all images in column B. Scalebar in 1E = 10 µm for all images in columns C-E. **F-H**. Measurements made on 15 neurons of the AX1D group, which includes 4 neurons with a 2-peaked 2°, 6 with shoulder+peak and 5 with a single-peaked 2°. See text for measurement details.

While some neurons had a single peaked 2° and others had two clearly apparent peaks, a subset of neurons displayed a shoulder melding into a peak (Fig. 5A), indicating that AIS and soma AP generation zones were more closely localized. To quantify this phenomenon, we calculated the slope of the 2° from the AIS peak (or shoulder) to the midpoint between the peaks (Fig. 5B). For all sham and most IN2D neurons, this slope was negative, allowing the peaks to be unambiguously distinguished. This measure could only be made on neurons whose 2° manifested either 2 peaks or a shoulder and a peak; consequently, a subset of neurons from the AX1D and AX2D groups with a single peak did not qualify. Eight of the 23 neurons within the AX1D group had positive slopes (while 5 had a single peak, Fig 6A). Normalizing the 2° to the second (or only) peak and averaging the waveform for all neurons in this group after aligning them to the time of the peak thus produced a relatively flat region of the curve where the expected AIS peak would occur (Fig. 6Bc). By the second day after injury, the axotomized group showed a smaller subset with either 1 peak or shoulder+peak (Fig. 6A), resulting in greater variability in the normalized AIS peak amplitude for the averaged waveform of this group (Fig. 6Bd). Surprisingly, the averaged waveform for the intact group at 1 day after injury appeared similar to that of the axotomized group 1 day after injury (Fig. 6B) due to 35% of this group manifesting the shoulder+peak profile (light blue symbols in Fig. 5B and also see Fig. 6A). For groups AX1D, AX2D, and IN1D, significantly fewer neurons had 2 peaks with a negative intervening slope, compared to the corresponding shams (z-tests, p<0.05, Fig. 6A).

**Figure 5.**
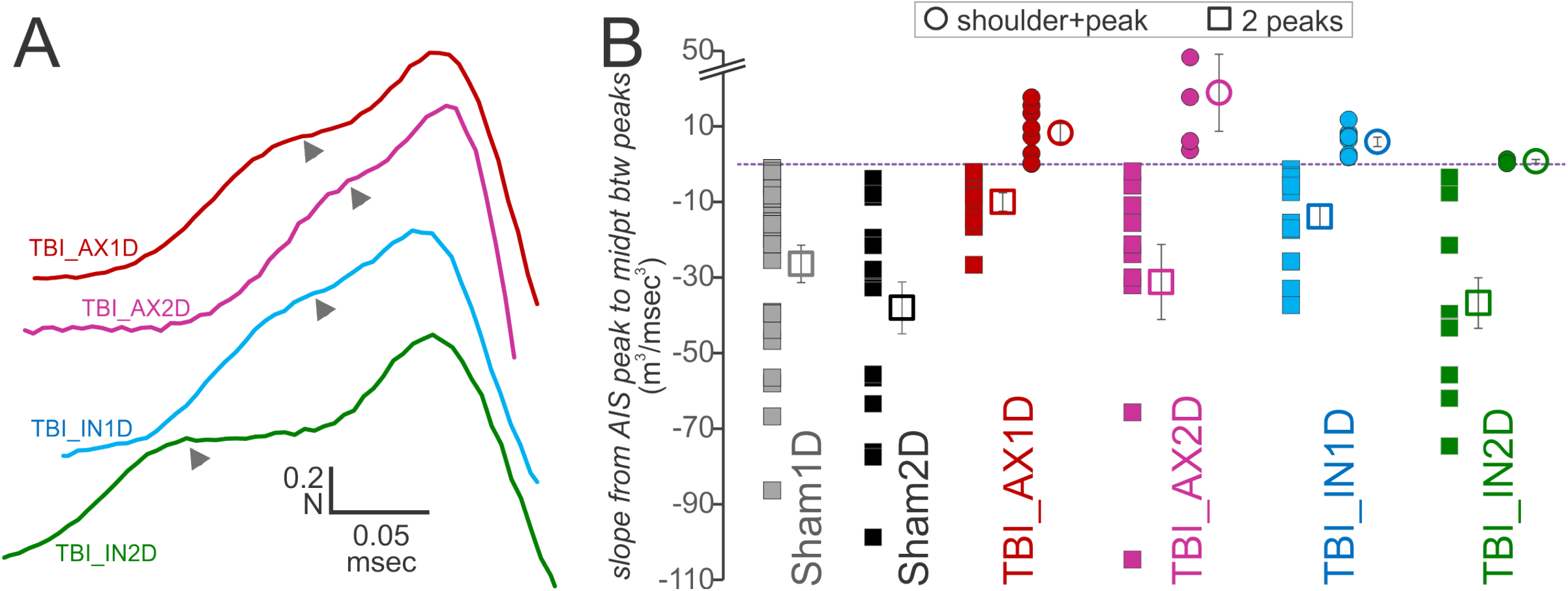
Some 2° waveforms manifest a shoulder prior to a peak. **A**. Examples of 2°s with a shoulder (arrowhead) prior to the peak. **B**. Slope of a linear line from the first peak to the midpoint between peaks is an effective measure to identify these shoulder neurons. Colors indicate the subject group. The 2° with 2 peaks are shown as squares, while those with a shoulder + peak are shown as circles. All of the cells with a shoulder had positive slopes, while all of the non-shoulder cells had negative slopes. Filled circles are from individual neurons, while empty circles shown mean + SEM. Note this measure cannot be made on cells with a single peak. n = 23 sham1D; 17 sham2D; 10 and 8 for AX1D with 2 peaks and shoulder+peak, respectively; 10 and 4 for AX2D with 2 peaks and shoulder+peak, respectively; 15 and 8 for IN1D with 2 peaks and shoulder+peak, respectively; and 12 and 2 for IN2D with 2 peaks and shoulder+peak, respectively.

**Figure 6.**
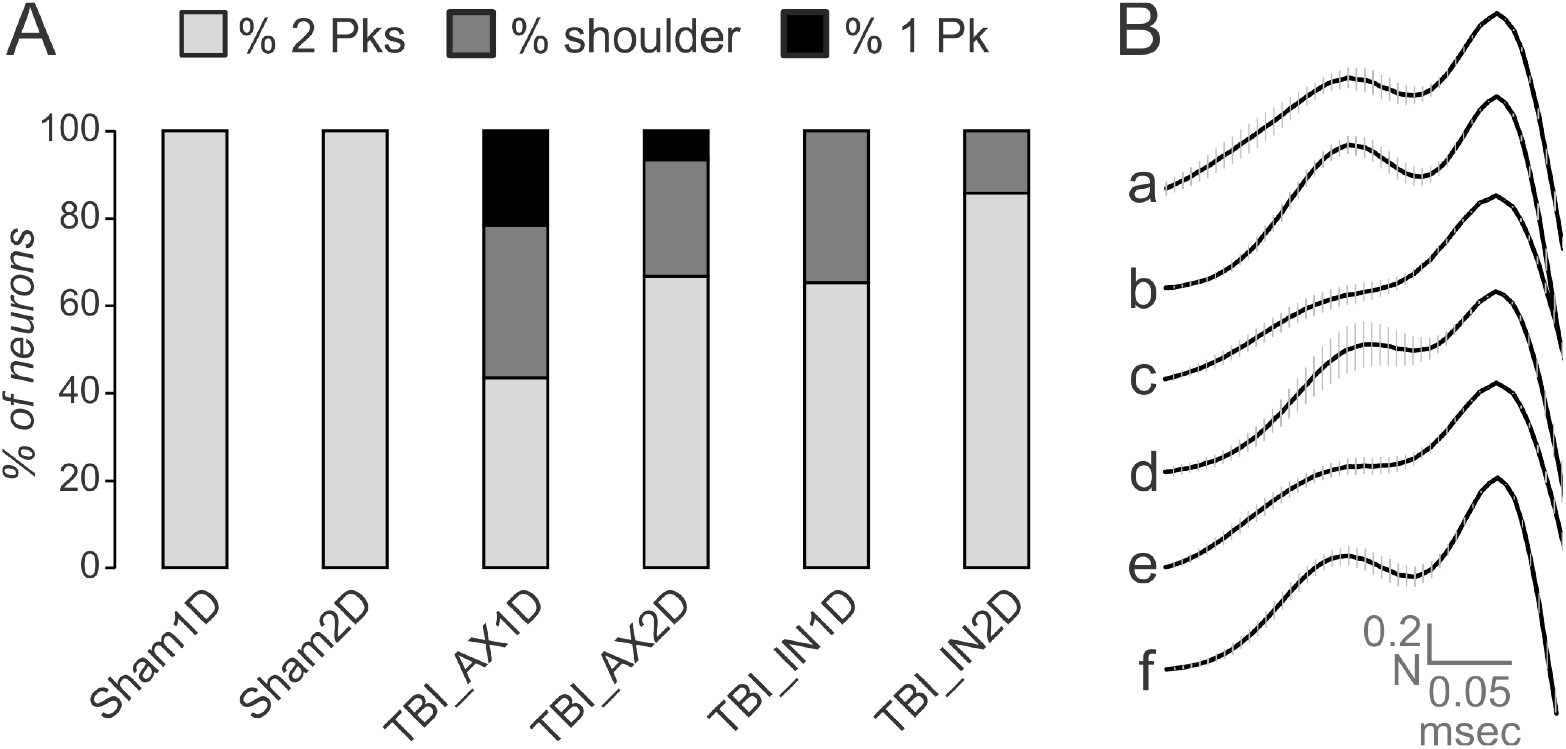
**A**. Proportion of neurons having 2, shoulder+1, or 1 peak(s) in the 2° of the AP. N = 23, 17, 23, 15, 23, and 14 cells for the groups as shown. **B**. The 2°s for all APs from a cell were normalized to the 2nd (or only) peak, aligned at the time of that peak and then averaged. The average was then taken for each subject group. The vertical lines indicate the SEM of this mean. Subject groups are: Sham1D (a), Sham2D (b), AX1D (c), AX2D (d), IN1D (e), IN2D (f). Z-tests showed that the TBI_AX1D and TBI_IN1D had significantly fewer 2-peaked 2o neurons compared to Sham1D, and that the TBI_AX2D had significantly fewer 2-peaked 2° neurons than Sham2D.

The timing of the single peak for the subset of neurons manifesting one peak within the AX1D and AX2D groups was examined to determine whether it corresponded to the AIS peak or the soma peak of 2-peaked 2° (Fig. 7). All first peaks from 2-peaked 2° (filled diamonds) and all shoulders from shoulder+peak 2° (open diamonds) occurred 0.15 msec or less after AP threshold. In fact, all but two occurred less than 0.14 msec after AP threshold (one in AX2D at 0.15 msec and one in IN2D at 0.144 msec). Most of the second peaks for 2-peaked 2° (filled squares) and shoulder+peak 2° (empty squares) occurred more than 0.15 msec after the AP threshold. Within the IN1D group, however, several occurred slightly before this time (see open light blue circles in Fig. 6A). The timing of single peaked 2° (triangles in Fig. 7) was most consistent with the timing of the second (somatic) peak; in fact, the timing of the single peak times was even later than that of the second peak in instances of 2 peaks. To examine this phenomenon further, the time of the second or only peak was compared across neurons with 2 peaks, a shoulder+peak or a single peak for the AX1D group. A one-way ANOVA showed a significant effect of peak number on relative timing, and the Bonferroni post-hoc showed that the time of the peak for single peaked neurons was significantly greater than that of the second component of the 2-peaked and the shoulder+peak neurons, suggesting that in the absence of an AIS-generated AP, more time is required to discharge the soma.

**Figure 7.**
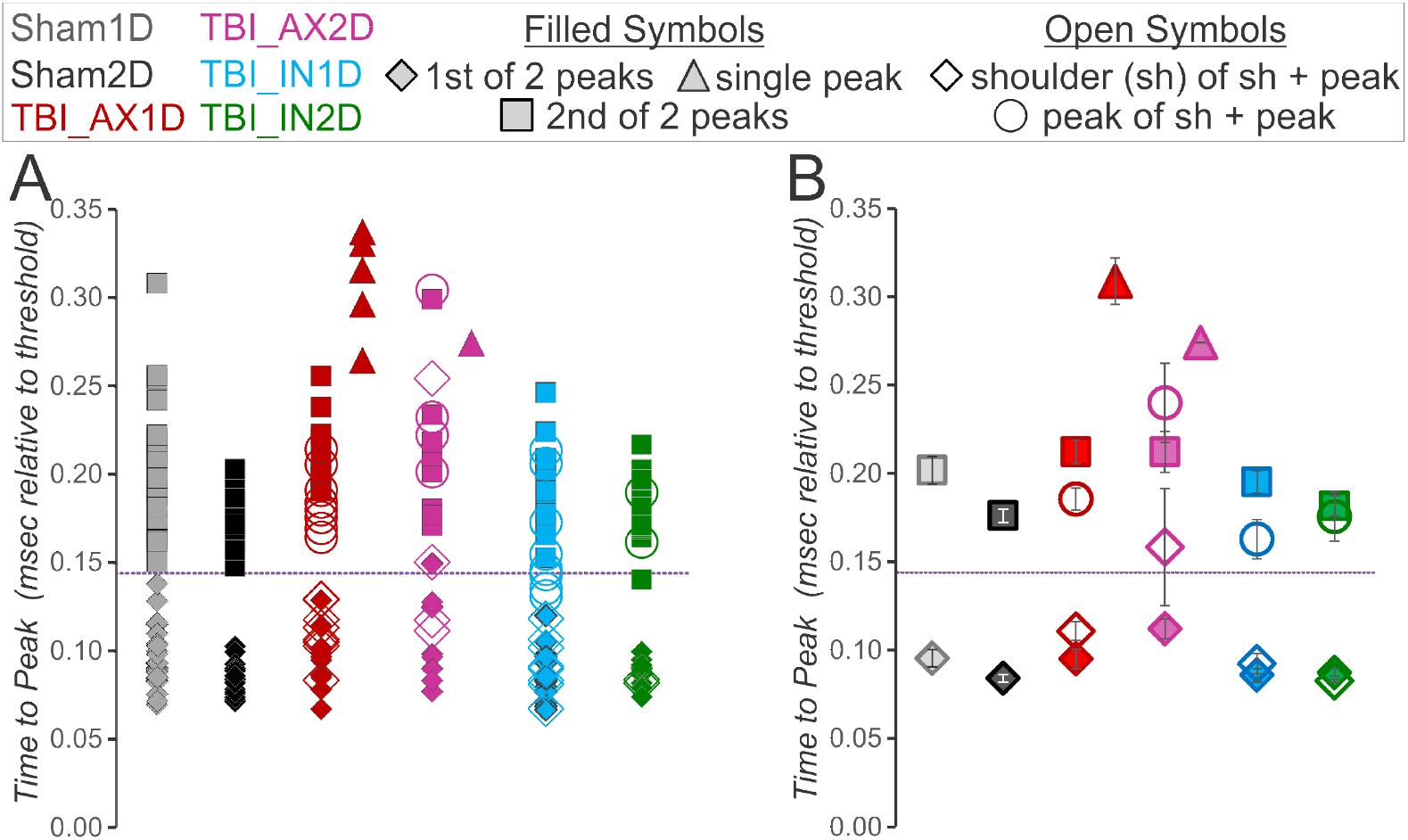
Most AIS or first peaks of the 2° occur less than 0.144 msec after threshold (dashed line), while most somatic or 2nd 2° peaks occur later than this. The single peaked 2° occur as late as or later than the soma time to peak. Time to peak for individual cells shown in **A** and means ±SEM shown in **B**. Color indicates subject group, as labeled. The AIS peaks are shown as diamonds, filled for 2-peaked and open for shoulder+peak 2°. The somatic peak is shown as squares, filled for 2-peaked and open for shouler+peak 2°. Single peaked 2° are triangles. N = 23 sham1D; 17 sham2D; for TBI_AX1D: 10 two-peaked, 8 shoulder+peak, 5 single-peaked; for TBI-_AX2D: 10 two-peaked, 4 sh+peak, and 1 single-peaked; for TBI_IN1D: 15 two-peaked, and 8 sh+peak; and for TBI_IN2D: 12 two-peaked and 2 shoulder+peak.

Our new measure of LS_diff_ differentiates single peaked and double peaked 2° waveforms (see Fig. 1). This measure is shown for experimental groups in Fig. 8, where once again one versus two peaked 2° could be differentiated at the threshold line of 2.25. As expected from our results with TTX application to the AIS, all 2° manifesting a single peak (triangles) fell below the threshold of 2.25, while all 2° with 2 peaks (squares) were above this threshold (Fig. 8). For the neurons that had a 2° with a shoulder+peak (circles), many had LS_diff_ values that fell below the threshold, as the 1 peaked 2°, while others were just above this threshold. A 2-way ANOVA found no significant difference between subject groups, but p<0.001 for the number of peaks, where shoulder+peak was considered 1.5 peaks. There was also no significant interaction. A Bonferroni post-hoc showed that LS_diff_ for 2-peaked 2° was significantly different from both the 1-peaked and the shoulder+peak 2°.

**Figure 8.**
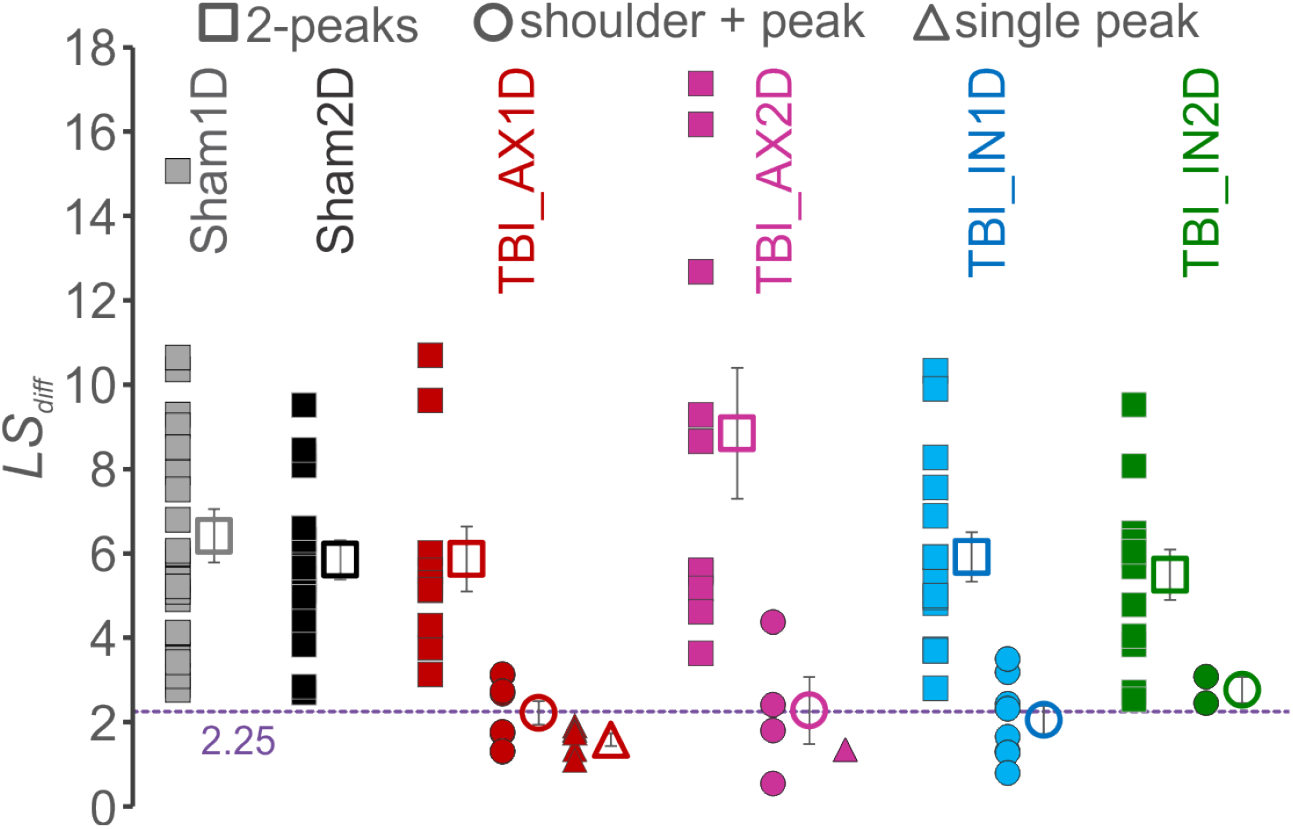
Measurement that differentiates between single and 2-peaked 2^o.^ The sum of the difference of actual values from the linear slope calculated from 99% of the second or only peak to 30% of that peak (LS_diff_). Individual neurons in solid symbols and mean ±SEM in open symbols. All 2-peaked 2° fall above the 2.25 line (purple dashed line), and all single-peaked 2° fall below this line. Shoulder+peak neurons tend to straddle the line. Number of cells are the same as for previous figure.

In order to examine the effects of concussive brain trauma on the relative voltage acceleration during the AP upstroke at the AIS and the soma, the peak amplitudes of the 2° for AIS and somatic locations were examined. The 2° peak amplitude at the AIS was significantly lower than sham for both AX1D and IN1D groups (Fig. 9A, ANOVA, Bonferroni post-hoc, p<0.05). The 2° peak amplitude at the soma was also significantly lower than sham for both AX1D and IN1D groups, as well as for the AX2D group (Fig. 9B, ANOVA, Bonferroni post-hoc, p<0.05).

**Figure 9.**
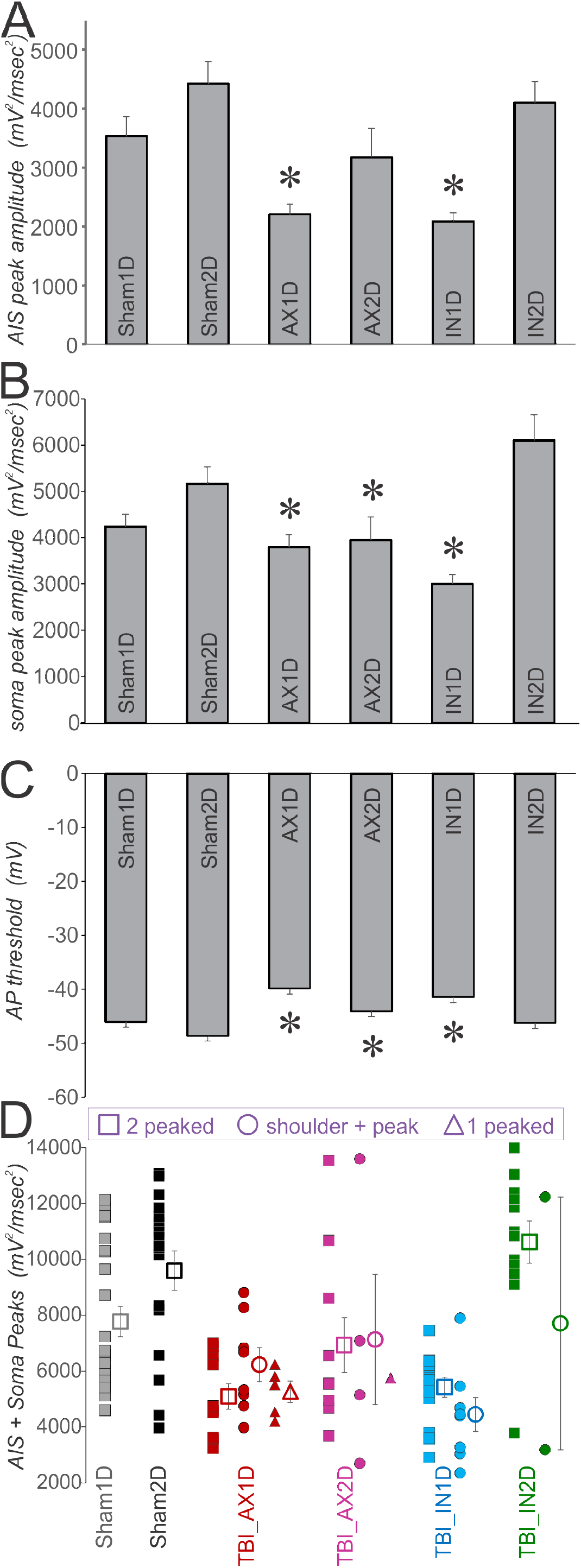
The 2° peak values suggest that AP generation at both the AIS and the soma are altered for some experimental groups. **A**. AIS 2° peak amplitude. N = 15, 13, 5, 4, 15, and 14 for groups as shown left to right. Note numbers of neurons are smaller for AX groups, since some APs had a single peak that was measured with the soma peak. * = ANOVA, with Bonferroni posthoc, p<0.05. **B**. Soma 2° peak amplitude. N = 23, 17, 18, 14, 23, 14 for groups as shown left to right. * indicates that these groups are significantly different from all 3 other groups (ANOVA, Bonferroni posthoc, p<0.05). **C**. Action potential threshold was more depolarized in the AX groups at 1 and 2 days and in the IN1D group compared to the shams and IN2D group. Numbers of neurons are the same as in B above. * indicates that these groups are significantly different from all 3 other groups (ANOVA, Bonferroni posthoc, p<0.05). **D**. The sum of AIS and Soma peaks shown per neuron. Neurons with two peaked 2° shown as squares, neurons with a 2° containing a shoulder and peak are shown as circles and those with a single peak are shown as triangles. Mean ±SEM shown in open symbols with error bars. Note that in the TBI_AX1D group, the 1 peaked 2° were similar in amplitude to the sum of both peaks for the 2 peaked 2°.

This difference returned to sham levels for the IN2D group. The three TBI groups showing reduced soma-regional AP acceleration also had a significantly more depolarized AP threshold that returned to near-sham levels for the IN2D group (Fig. 9C, ANOVA, Bonferroni post-hoc, p<0.05).When the sum of the AIS-and soma-regional 2° peak amplitudes was calculated, presumably reflecting the total sodium current density during the AP upstroke, the sums for the AX1D, AX2D and IN1D groups were significantly less than sham and IN2D groups (Fig. 9D, ANOVA, Bonferroni post-hoc, p<0.05). However, we also observed that the amplitude of 1-peaked 2° was similar to that of the sum of the two peaks of 2-peaked 2° within the AX1D group. This result suggested that the AIS and somatic generation zones may approach each other more closely. If so, it follows that the peak amplitude of the 1-peaked 2° of the AX1D group should be larger than that of the somatic peak of the 2-peaked and shoulder+peak 2° in the AX1D group. We find this result to be the case (Fig. 10A, ANOVA, Bonferroni post-hoc, P<0.05).

**Figure 10.**
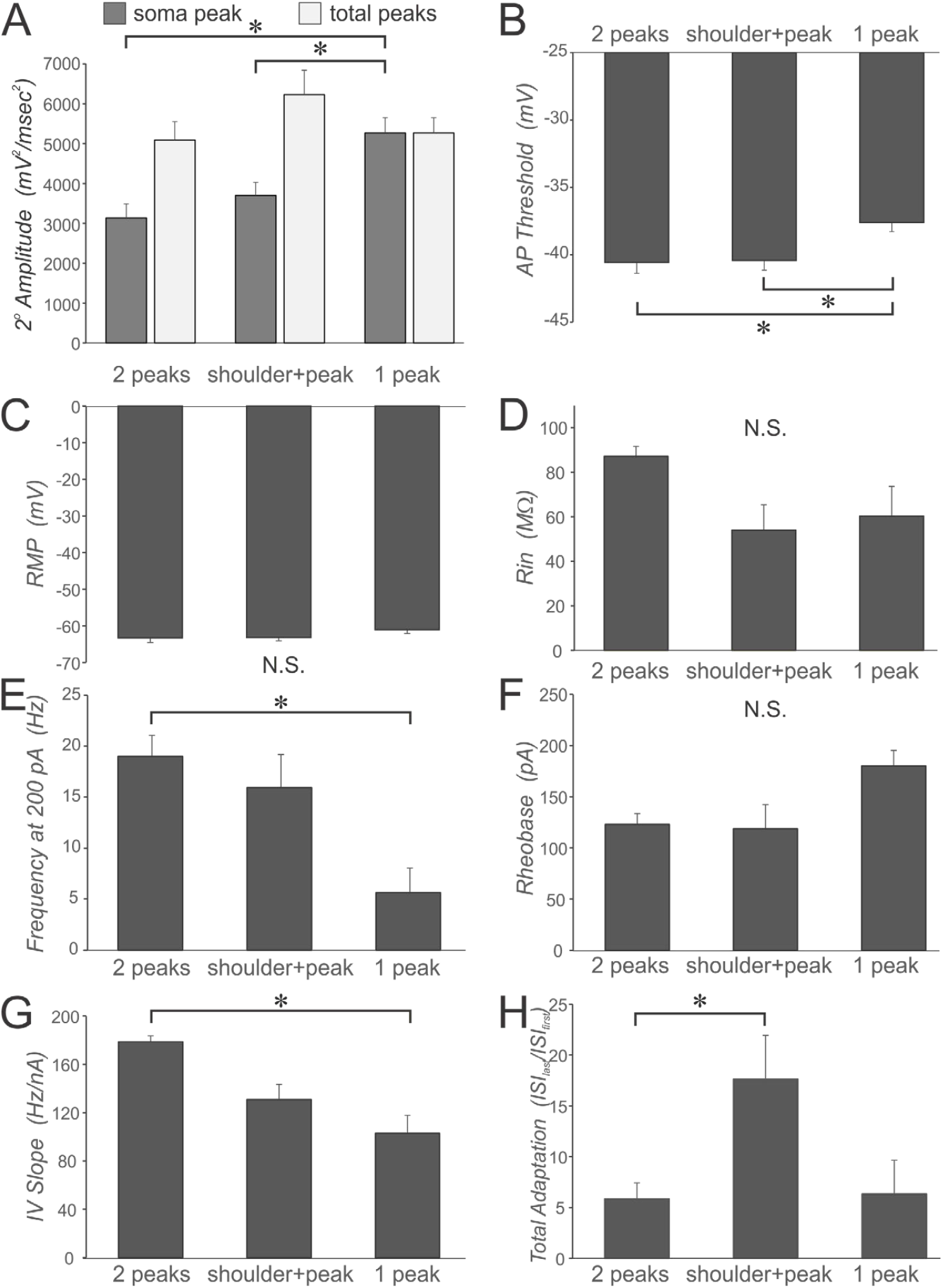
Effect of 2° peak number and shape for neurons from the AX1D group. In all cases a 1-way ANOVA with Bonferroni post-hoc was performed, *= p<0.05, with 10 2-peaked, 8 shoulder + peak and 5 1-peaked neurons. **A**. The 2° amplitude of the somatic (or only) peak is shown in gray, and the combined peak amplitude of the AIS and somatic (or only) peak(s) is shown in white. The total 2° peak amplitude is not different for 1-peaked neurons compared to that of 2-peaked neurons or compared to those with a shoulder+peak. The somatic peak alone is significantly higher for the 1-peaked neurons compared to the other groups. This result suggests that the AIS generation site may have moved towards the somatic generation site such that the two are no longer differentiated, or that abolition of AIS function gives rise to a homeostatic upregulation of somatic sodium channels. **B**. The threshold for AP generation was significantly different for the 1-peaked neurons compared to the groups. **C & D**. There were no significant differences in resting membrane potential or input resistance among these groups, suggesting that 1-peaked neurons are healthy by conventional criteria. **E**. The frequency of APs generated during a 400 msec depolarizing pulse at 200 pA was significantly compared less in the 1-peaked to 2-peaked neurons. **F**. Rheobase, or the lowest depolarizing current to generate an AP was not significantly different for these groups. **G**. The slope of the plot of AP frequency versus injected current level was significantly lower for 1-peaked neurons compared to 2-peaked neurons. **H**. The total adaptation was calculated as the interval between the last two APs divided by the interval between the first two APs during depolarizing current injection. Shown here is the maximum total adaptation across all sweeps. The shoulder+peak group manifested significantly greater adaptation than the 2-peaked group.

The impairment of AIS-regional AP generation in the AX1D, AX2D and IN1D groups did not arise as a function of unhealthy neurons as judged by conventional criteria. The input resistance in these groups was similar to that of the sham and IN2D groups (Table 1), although the resting membrane potential (RMP) was slightly more depolarized in these groups (Table 1). In addition, the 1-peaked or shoulder+peak neurons within the AX1D group were not significantly different from the 2-peaked AX1D neurons on resting membrane potential and input resistance (Fig. 10C, D), suggesting the differences in the shape of the 2° cannot be explained by differences in global metrics of neuronal health.

We next sought to correlate the consequences of trauma-induced AIS-regional functional perturbation to the associated AP discharge patterns of affected neurons. We observed differences in the AP discharge patterns of 1-peaked and shoulder+peak neurons compared to neurons manifesting 2 peaks. In particular, the AP threshold for 1-peaked AX1D neurons was significantly depolarized compared to either shoulder+peak or 2-peaked neurons (Fig. 10B), yet the rheobase was not significantly different between these three AX1D groups. The 1-peaked neurons discharged APs at a significantly lower frequency in response to 200 pA depolarizing current injection, and the overall slope of the plot of AP frequency versus current injection was significantly lower compared to the 2-peaked neurons (Fig. 10 E, G). These results suggest that the 1-peaked neurons are less excitable.

The total adaptation across a train of APs was measured as the last inter-spike interval divided by the first inter-spike interval. We compared the maximum total adaptation and found that the shoulder+peak neurons demonstrated significantly more adaptation than 2-peaked neurons.

We previously found that aberrations of intrinsic cellular and membrane properties were prevented or ameliorated when experiments were conducted on brains extracted from CypDKO mice subjected to the equivalent concussive TBI (Sun & Jacobs, 2016). Here, we examined whether CypDKO might mitigate trauma-induced effects on AIS-and soma-regional voltage acceleration. Within CypDKO mice, we found that even when axotomy occurred just proximal to the soma (not shown), the AP 2° still manifested 2 peaks, or in a few cases, a shoulder+peak (Fig. 11). Single-peaked 2° were not observed in any neurons from CypDKO mice. In fact, the overall shape of the 2° was fairly similar in axotomized and intact groups compared to the shams (Fig. 11B). The measure allowing identification of the shoulder+peak 2° neurons reveals only 4 neurons with positive slopes measured from the AIS peak to the midpoint between the AIS and somatic peaks (Fig. 12A). Among those 4 shoulder+peak CypDKO neurons, only 2 fell just below the 2.25 threshold on the LS_diff_ measure (Fig. 12B). Thus, CypDKO was associated with a striking amelioration of trauma-induced perturbation of AP upstroke kinetics (compare Figs: 5B and 12A; 6 and 11; 8 and 12B).

**Figure 11.**
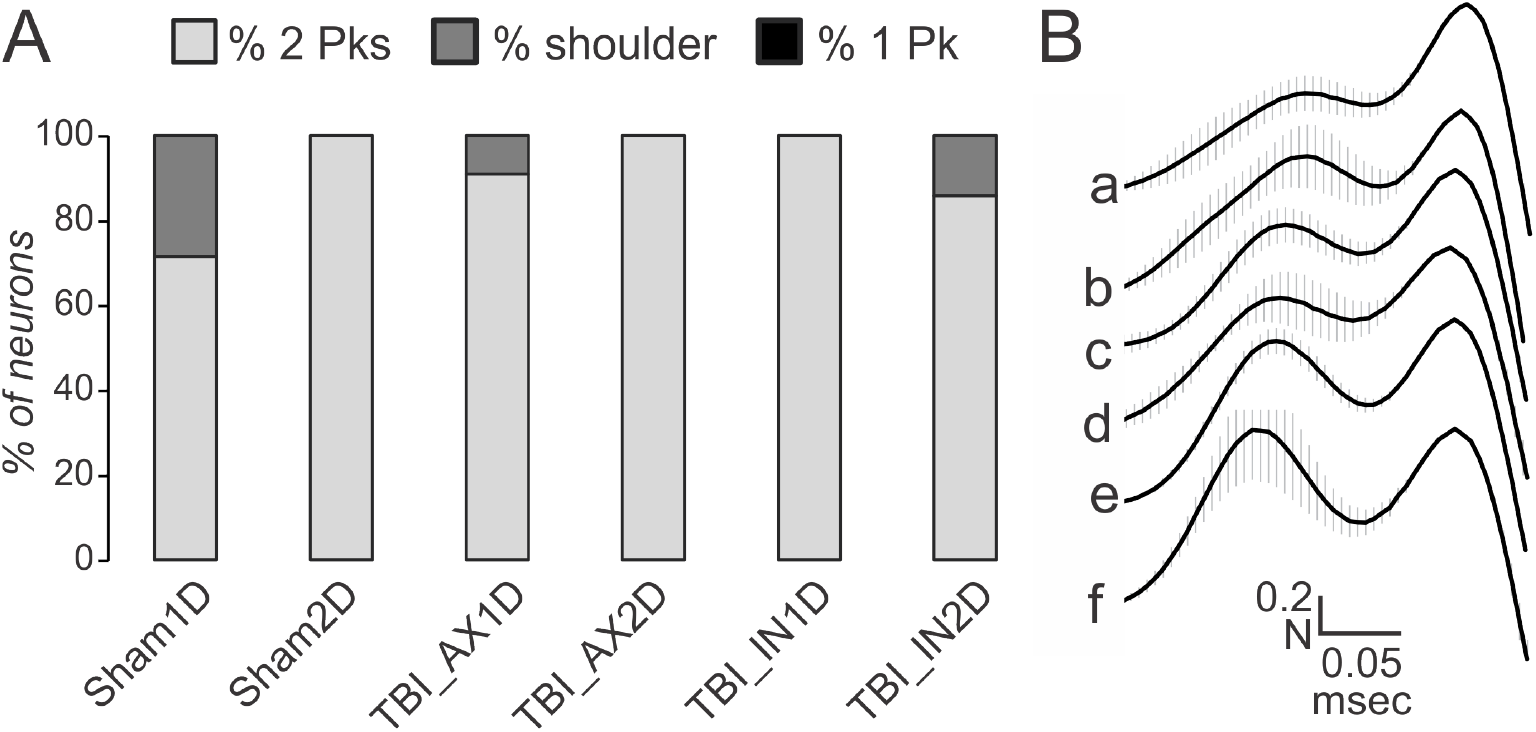
Effects of mTBI on the morphology of 2o waveforms in CypDKO tissue. **A**. For neurons recorded in slices from CypDKO animals, proportion of neurons having 2, shoulder+1, or 1 peak(s) in the 2° of the AP. n = 7, 8, 11, 6, 13, and 7, for the groups as shown. **B**. The 2°s for all APs from a cell were normalized to the second (or only) peak, aligned at the soma Peak of the 2°, and then averaged. For each subject group, the average of these cell averages was then taken, with the vertical lines indicating the SEM. Subject groups are: CypDKO_Sham1D (a), CypDKO_Sham2D (b), CypDKO_AX1D (c), CypDKO_AX2D (d), CypDKO_IN1D (e), CypDKO_IN2D (f). Numbers of neurons averaged are the same as indicated for A.

**Figure 12.**
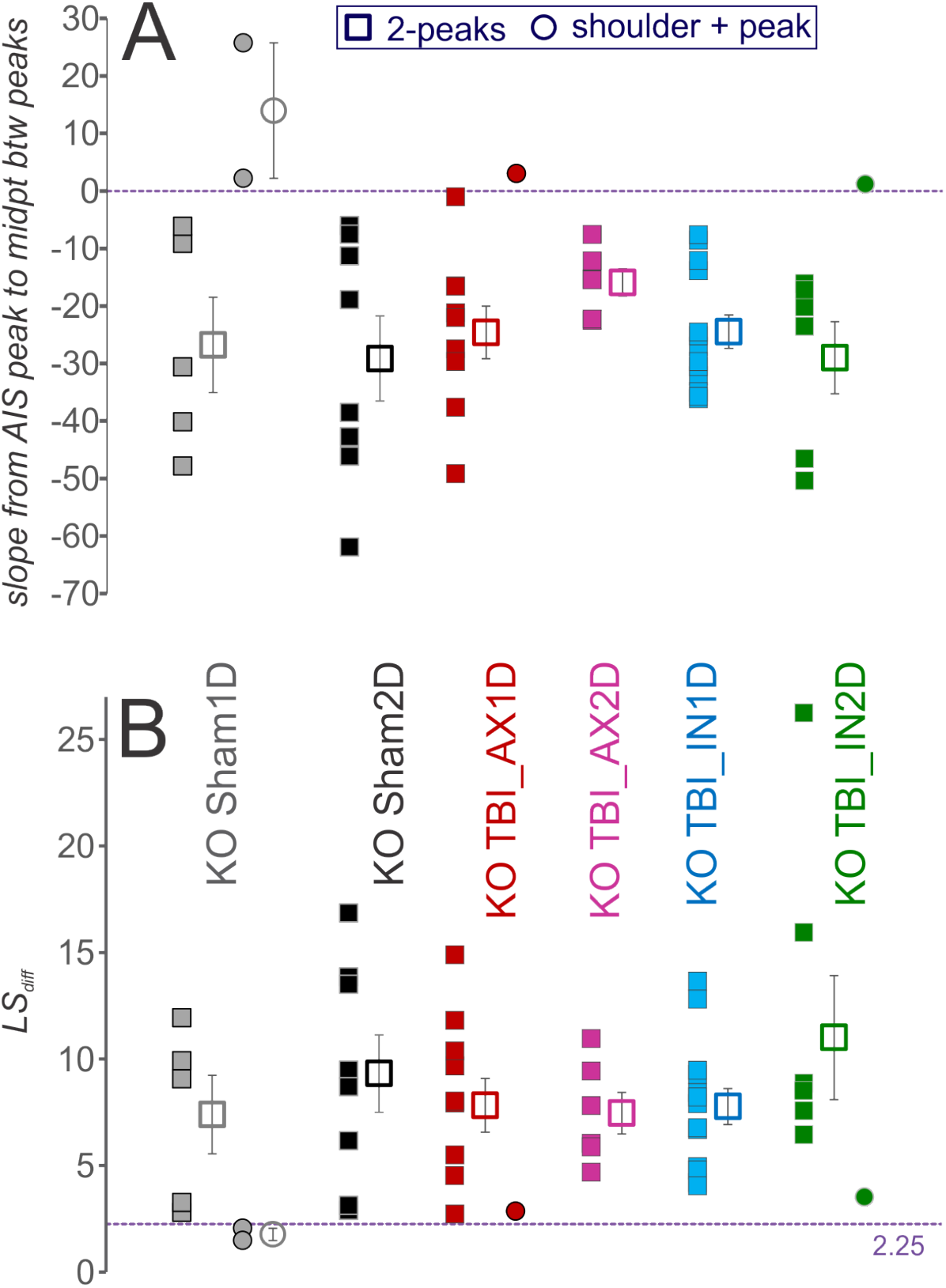
Effect of CDKO on the shape of the 2°. **A**. Slope from the AIS peak to the midpoint between AIS & soma peaks. Means ±SEM shown in open symbols with error bars. All but 4 of the CDKO neurons’ 2° had negative slopes. **B**. Summed difference from linear slope for CDKO groups, mean ±SEM shown in open symbols with error bars. Here only two neurons’ 2° had a summed difference from linear slope below the threshold of 2.25. For A and B, CDKO 2-peaked 2°, n = 7, 8, 9, 6, 13, and 6 neurons for groups as shown.

In order to compare further between WT and CypDKO on a series of measures (Fig. 13), 2-way ANOVAs with Bonferroni post-hoc tests were performed, with the main effects listed in Table 2. For the number of cells in each group, see Table 2. For the post-hoc interactions between subject group and the experimental conditions of WT or CypDKO, significance is shown in Fig. 13, with * denoting p<0.05.

**Table 2.** List of p values for the Main Effects of a 2-way ANOVA for all measures shown in figure 13. Letter to the left of measure name is consistent with that shown in figure 13. Significant effects of a 2-way ANOVA of subject group (sham1D, sham2D, AX1D, AX2D, IN1D, IN2D) and experimental condition (WT vs CypDKO) shown with * (p<0.05). Numbers of cells for each group: WT Sham1D 23; WT Sham2D 17; WT AX1D 23; WT AX2D 15; WT IN1D 23; WT IN2D 21; CypDKO Sham1D 8; CypDKO Sham2D 11; CypDKO AX1D 11; CypDKO AX2D 6; CypDKO IN1D 13; CypDKO IN2D 7.

|  | **Experimental Condition** | **Subject Groups** | **Interaction** |
| --- | --- | --- | --- |
| **A AIS peak amplitude** | 0.351 | <0.0001* | 0.003* |
| **B soma peak amplitude** | 0.004* | <0.0001* | <0.0001* |
| **C time to AIS peak** | 0.465 | 0.0001* | 0.022* |
| **D time to soma peak** | 0.041* | 0.014* | 0.007* |
| **E AP threshold** | 0.775 | 0.004* | <0.0001* |
| **F % Bumpiness** | 0.177 | 0.143 | 0.045* |
| **G LS<sub>diff</sub>** | <0.0001* | 0.132 | 0.064 |
| **H time less than AIS peak** | 0.0001* | 0.767 | 0.167 |

**Figure 13.**
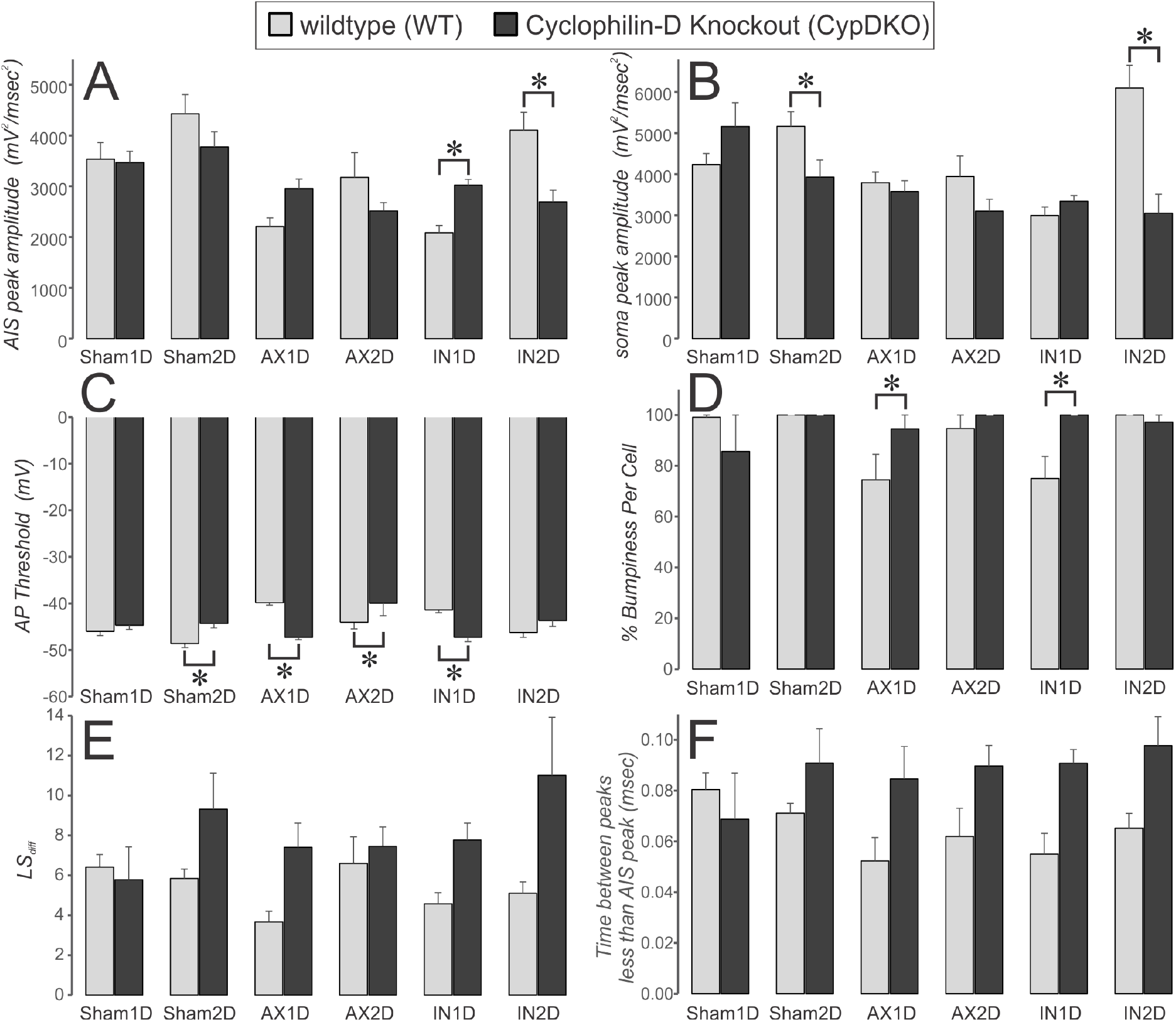
Comparison between WT and CypDKO experimental conditions for all subject groups. WT is in lighter gray and CypDKO is in darker gray. Subject groups are marked at the bottom of the columns. For each measure, a 2-way ANOVA was performed. See Table 2 for p values for subject group, experimental condition and interactions. When a significant interaction between subject group and experimental condition was present, a Bonferroni post-hoc analysis was performed, with significant effects indicated here by an *.

For the AIS peak amplitude there was a significant effect of subject group, with AX1D and IN1D being significantly lower than Sham1D. The AIS peak for AX2D as well as AX1D and IN1D was significantly lower than Sham2D. For the IN2D group, the AIS peak was significantly different from AX1D and IN1D. The interaction between subject group and experimental condition was also significant, with the post-hoc test showing the following. The reduced 2° for the AIS peak was recovered in the CypDKO mice only for the IN1D group. Surprisingly, while the AIS peak was increased in CypDKO compared to WT for the IN1D neurons, it was significantly decreased in CypDKO compared to WT for the IN2D neurons. It appears that restricting the assembly of the mitochondrial pore transition complex within CypDKO mice may be associated with nonlinear effects on neuronal excitability (Sun & Jacobs, 2016).

For the AP acceleration at the soma (Fig. 13B), there was a significant effect of subject group, experimental condition and the interaction between these two. The IN1D group had a significantly smaller soma-regional peak 2° compared to the Sham1D and Sham2D groups. The IN2D group was significantly different from AX1D, AX2D and IN1D groups. For the interaction between subject group and experimental condition, surprisingly there was an effect on the Sham2D neurons, with a reduction in the somatic 2° peak in the CypDKO condition compared to WT. There was also a reduction in this peak under CypDKO condition for the IN2D group.

For the AP threshold (Fig. 13C), there was a significant effect of subject group, no effect of experimental condition, but a significant interaction of subject group and experimental condition. The AX2D group had a significantly more depolarized AP threshold compared to the Sham2D group. For the interaction, the CypDKO condition ameliorated the more depolarized AP threshold for both AX1D and IN1D groups but exacerbated it in the AX2D group and produced a more depolarized AP threshold in the Sham2D group. The overall result of the CypDKO condition was a very similar AP threshold level between subject groups.

In previous examinations of the 2° of the AP upstroke, authors have characterized the shoulder+peak 2° waveform as possessing a range of *bumpiness*, depending on whether the slope of the 2° passes through zero (Kress *et al*., 2008). The phenomenon of less bumpiness was attributed to a closer relative location of the AIS and somatic AP initiation zones (Kress *et al*., 2008). Here we made this measurement explicitly for the 2° of each AP, then calculated the % of APs per cell that were bumpy, which we defined as any 2° waveform in which the intervening slope between AIS and somatic peaks passed through zero. This measure was then averaged among neurons of each subject groups (Fig. 13D). For some groups, every single 2° slope passed through zero, thus putting the average at 100% without error (WT: Sham1D, Sham2D, and IN2D; CypDKO: Sham2D, AX2D, and IN1D). The three experimental groups of the WT condition most affected by TBI (AX1D, AX2D and IN1D) all contained some non-bumpy 2° neurons. The 2-way ANOVA showed no significant effect of subject group nor of experimental condition, but did show an interaction between these two, with a significant increase in bumpiness for both AX1D and IN1D groups. Thus, within these groups the CypDKO condition gave rise to enhanced bumpiness, presumably due to a greater spatial separation between AIS and soma.

While the bumpiness quotient can only be measured in 2° with 2 peaks, the LS_diff_ measure can determine how close the 2° is to a single peak for all 2°. This metric provided a more inclusive measurement of bumpiness (Fig. 13E). We found a threshold level at 2.25, where lower values were typical for 1-peaked 2°, and higher values were typical for 2-peaked 2° (Figs. 1F, 8). A 1-way ANOVA with Bonferroni post-hoc for the WT condition showed a significantly lower LS_diff_ for the AX1D group compared to the Sham1D. This result was ameliorated in the CypDKO condition. The 2-way ANOVA considering both experimental conditions showed no significant effect of subject group but a significant effect of experimental condition, with an increase in LS_diff_ under the condition of CypDKO. There was a trend towards an interaction between subject group and experimental condition.

The examination of the CypDKO 2° suggested that under this condition, there were more 2° with 2 peaks and that the minimum between the peaks was lower than for the WT condition. We evaluated this by measuring the duration of time between the two peaks (only for 2-peaked or shoulder+peak 2°) during which the value was less than that of the AIS peak (Fig. 13F). Here, there was a significant effect of experimental condition. Relative to the WT condition, CypDKO produced 2° with greater durations of time during which the 2° was less than that of the AIS peak. This result suggests that the current producing the AP at the AIS decays to a greater degree prior to the generation of the current producing the AP at the soma. Thus, this result supports the bumpiness quotient result in suggesting that the AIS and soma AP initiation zones are further apart in the CypDKO condition compared to WT.

## Discussion

The results of the present study represent an advancement in the use of a detailed, quantitative analysis of AP upstroke kinetics as an assay of the functional integrity of the neuronal AIS. Although the present study exploited this assay in the context of a well-characterized model of concussive TBI, we propose its broad utility to shed light on the pathobiological mechanisms underpinning neurological disease in general (Favero *et al*., 2018). While past research had implicated the AIS as a locus of vulnerability to traumatic axotomy, the physiological consequences of this phenomenon remained unclear. By means of analysis of the 2° with respect to time of the membrane voltage during the AP upstroke within a subset of layer 5 pyramidal neurons, we were able to disentangle and rigorously quantify the relative contribution of the AIS and the soma to the AP upstroke. Via this analysis, we discovered a subset of axotomized layer 5 pyramidal neurons which manifested a single peak corresponding in time to the second peak of neurons with two peaks, a result consistent with trauma-induced abolition of AIS function in these neurons. Our LS_diff_ measure distinguished this 1-peaked group of neurons from the 2-peaked group (Fig. 8). Moreover, we determined that there exists a continuum of trauma-induced, AIS-regional dysfunction, in which outright abolition of the AIS-regional peak represents one extreme, while the manifestation of two clearly appreciable peaks separated by a negative intervening slope represents the other. Bounded by these extremes exists a sub-set of both morphologically intact and axotomized neurons that manifest a biphasic waveform in which the AIS-regional portion melds into the soma-regional portion, a waveform we have classified as *shoulder+peak* (Fig. 5).

Post-hoc measurements of biocytin-filled, axotomized YFP^+^ layer 5 pyramidal neurons stained for the AIS scaffolding protein ankyrin-G revealed a lack of definite correlation between the site of emergence of axonal retraction bulbs and AIS-regional functional perturbation, as some neurons with bulbs at the AIS nevertheless manifested dual-peaked 2° waveforms (Fig. 4D_4_). However, there existed a subset of axotomized neurons 1 and 2 days after injury in which the AIS appeared severely maimed, assuming a club-shaped morphology of increased width (Fig. 4G); further, these neurons displayed filipodia-like axonal sprouts that may reflect an attempt at post-traumatic axonal regeneration. This morphology corresponded to all axotomized neurons manifesting abolition of the AIS-regional contribution to the AP upstroke upon 2° analysis. Among the subset of axotomized neurons, one day after injury, abolition of AIS-regional contribution to the AP upstroke correlated with electrophysiological perturbation across several metrics. Such neurons displayed a significantly depolarized AP threshold, a significantly reduced firing frequency at 200 pA, and a significantly lower F-I slope compared to dual-peaked and shoulder+peak counterparts (Fig. 10). Thus, this type of axotomy renders neurons significantly less excitable.

The lack of definite correlation between the site of emergence of axonal retraction bulbs and corresponding AIS functional perturbation may indicate that heterogeneity within the intracellular signaling cascades set into motion by concussive trauma drives the functional consequences at the AIS. Consistent with this notion, previous work has indicated that vibratome-induced axotomy even in the vicinity of the AIS does not give rise to aberration of AIS-regional function (Palmer & Stuart, 2006). Thus, transection *per se* does not seem to drive AIS functional pathology, and neurons axotomized in this manner do not manifest any detectable aberration of intrinsic properties (Sun & Jacobs, 2017). Expanding upon this theme, here we show that restriction of the formation of the mitochondrial pore transition complex within CypDKO mice ameliorates many aspects of the trauma-induced AIS-regional functional perturbation in comparison to the Thy-1 YFP-H (WT) condition. In particular, the reduction of the AIS peak of the 2° in AX1D and IN1D (Fig. 9A) is partially ameliorated (Fig 13A). Interestingly, the decrease of the somatic peak for AX1D, AX2D and IN1D (Fig. 9B) is not improved in the CypDKO mice (Fig. 13B).

These results inform the ongoing theoretical question of the relation between neuronal excitability and the distance relative to the soma of AIS emergence (Goethals & Brette, 2020). Seminal work has demonstrated that global enhancement of excitability by administration of high extracellular [K^+^] reliably induces translocation of the AIS away from the soma and associated homeostatic decrease in excitability within hippocampal neurons *in vitro* (Grubb & Burrone, 2010; Wefelmeyer *et al*., 2015). However, the AIS emerges further from the soma in layer 5 pyramidal neurons with axon-bearing dendrites, and these neurons possess a corresponding hyperpolarized AP threshold compared to neurons with axons emerging from the soma (Hamada *et al*., 2016). Thus, the relation between AIS start position and excitability seems to be nonlinear, arising in dependence on a diversity of variables. One *in silico* study (Gulledge & Bravo, 2016) using simplified models of neurons of a range of sizes and elaborating a varying quantity of primary dendrites with and without spines implicated larger surface area and more dense dendritic spines as crucial variables. In particular, within larger neurons more densely studded by dendritic spines—consistent with the neurons under study in the present communication—distal translocation of the AIS was predicted to enhance excitability, a relation attributed to the influence of dendritic architecture on AIS-regional values of resistance and the membrane time constant. Further, increased electrotonic isolation of the AIS from the soma may support enhanced excitability by reducing the pool of inactivated voltage-gated sodium channels (Kole & Stuart, 2012). The depolarized AP threshold encountered in AX1D and IN1D groups (Fig. 9C) was ameliorated within the CypDKO, although not in the AX2D group (Fig. 13C). Our metric of *% bumpiness* may reflect the relative separation of the AIS and somatic activation zones (Kress *et al*., 2008). In the CypDKO mice, this measure more closely approached control levels for AX1D and IN1D groups (Fig. 13D). Similarly, the time between peaks during which the 2° is less than the AIS peak value may reflect the distance between the AIS and somatic activation zones, and that time was increased in the CypDKO mice (Fig. 13F). Within these same groups, a more hyperpolarized AP threshold was recorded in the CypDKO mice. Therefore, the increased separation of AIS and somatic activation zones as assessed by a surrogate measure correlates with enhanced excitability under these conditions.

Induction of ischemia via the middle cerebral artery occlusion model of ischemic stroke has been shown to give rise to the calpain-mediated cleavage of ankyrin-G and the degradation of the AIS (Schafer *et al*., 2009). Critically, this ischemia-induced disruption of AIS integrity led to the loss of neuronal polarity, the maintenance of which depends upon the presence of ankyrin-G (Hedstrom *et al*., 2008). Moreover, calpain-mediated proteolysis of voltage-gated sodium channels, including Na_V_1.2 (von Reyn *et al*., 2009) and Na_V_1.6 (Brocard *et al*., 2016), has been demonstrated in multiple models of traumatic injury to the nervous system. Thus, in the present study, abolition of the AIS-regional 2° peak within a sub-set of axotomized YFP^+^, layer 5 pyramidal neurons may arise secondary to calpain-induced proteolysis of ankyrin-G and consequent dissolution of the AIS and loss of neuronal polarity. It is conceivable that such neurons may attempt to mount a regenerative attempt by transforming a dendrite into an axon during the epoch of acute recovery after trauma (Gomis-Rüth *et al*., 2008). Alternatively, a reparative attempt could be mounted at the site of the dissolved AIS, thus explaining the emergence of sprouts from the stumps of the former location of the AIS (Fig 4C_1-2_). Because these pathobiological cascades arise in dependence on calcium, knocking out CypD may support the capacity of AIS-resident mitochondria to buffer the intracellular calcium that permeates neurons haphazardly secondary to trauma and thus support AIS functional integrity. Because AIS homeostatic plasticity in the form of morphological re-modeling itself arises in dependence on calcium (Evans *et al*., 2013), AIS dissolution secondary to neurological insult like ischemia and concussive trauma may represent the most extreme manifestation of this cell-intrinsic mechanism (Zhao *et al*., 2020).

Given the critical role of the AIS in tuning neuronal excitability toward the maintenance of a target range of output in the face of fluctuations in network excitatory-inhibitory balance, the effects of trauma-induced AIS dysfunction on neuronal plasticity represents a fascinating topic of future inquiry. Of particular interest are intact neurons, some minority of which we have shown manifest significant AIS-regional functional perturbation (Figs. 3, 5, 6 and 8). At 1 day after mTBI, we have demonstrated that based on intrinsic properties, relative synaptic input and, presently, based on AIS function, intact neurons are functionally similar to their axotomized counterparts (Greer *et al*., 2012; Hånell *et al*., 2015). Given their intact morphology, however, these neurons maintain their contribution to the dynamics of the neocortical network. The dysfunction of the AIS likely perturbs the tight correlation between neuronal input and consequent output—in other words, the precision of AP firing— (Lazarov *et al*., 2018), and such disruption during the post-traumatic epoch may underlie the altered sensory perception that can arise in patients after clinical TBI (Spiegel *et al*., 2016).

Might trauma disrupt the capability of these neurons to modulate excitability toward the maintenance of network homeostasis? Unlike axotomized counterparts, these neurons retain innervation of post-synaptic targets. If trauma can abrogate such a critical mechanism of plasticity, it is conceivable that previously manageable instances of stress could be rendered pathological within neuronal networks, at least during the first days following concussive insult. In support of the plausibility of this notion, traumatic brain injury—including the mild/concussive form—is a documented risk factor for the emergence of post-traumatic stress disorder (Iljazi *et al*., 2020). Finally, it must be considered that the impairment of AIS function especially within intact neurons may represent a neuroprotective mechanism to limit the spread of aberrant excitatory activity in the acute phase after trauma. Even with mild TBI, interictal-like activity can be observed early after injury (Hånell *et al*., 2015).

Notably, all YFP^+^ layer 5 pyramidal neurons targeted within the present study belonged to the Type A subclass of layer 5 pyramidal neurons. This emerging scheme of classification seeks to assign layer 5 pyramidal neurons to one of two subclasses (A or B) based on the presence of an I_h_-mediated depolarizing voltage sag in response to hyperpolarizing current injection and the presence of a depolarizing afterpotential upon cessation of hyperpolarizing current injection (Fig. S1) (Dembrow *et al*., 2010; Gee *et al*., 2012). Critically, these physiological metrics have been shown to correlate reliably to anatomical and morphological differences between these cardinal subtypes. In particular, type A neurons tend to elaborate wider and more geometrically complex apical dendritic arbors; to put forth extra-telencephalic axons; and to receive preferential inhibitory tone from both somatostatin-(Hilscher *et al*., 2017) and parvalbumin-expressing (Lee *et al*., 2014) inhibitory interneurons within the neocortex. This result augments past work demonstrating that virtually all YFP^+^ neurons within the Thy1-YFPH strain elaborate extra-telencephalic axons (Porrero *et al*., 2010). Thus, a potentially fruitful topic of future inquiry concerns the relative vulnerability to traumatic axotomy between type A and type B layer 5 pyramidal neurons. Might extra-telencephalic (type A) layer 5 pyramidal neurons be more susceptible to axotomy than their corticocortical (type B) counterparts? Such a difference, if present, might lead to novel insights concerning the cellular and physiological properties that predispose neurons to axonal injury.

## Supplement

**Figure S1.**
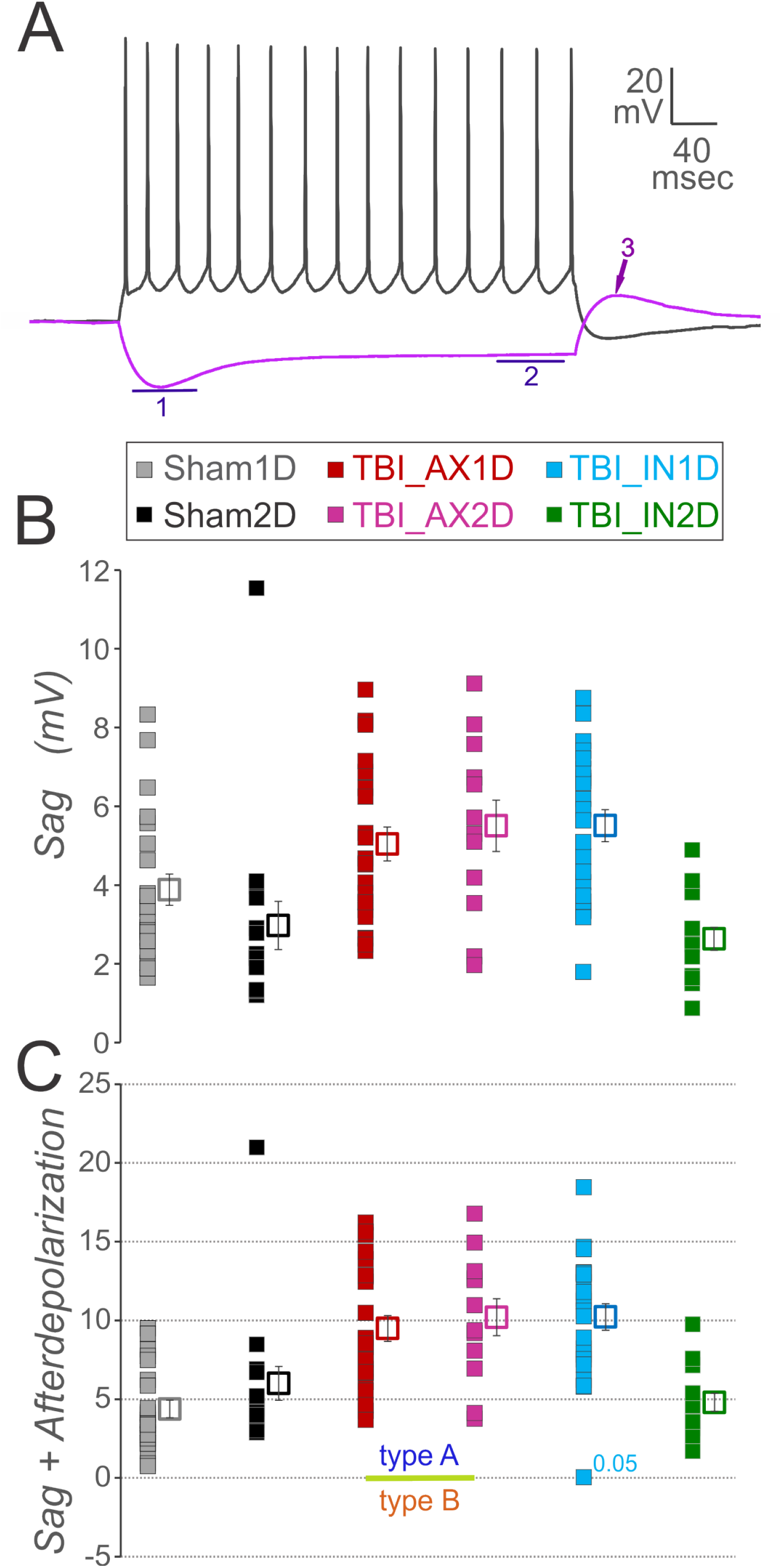
Layer 5 pyramidal neurons labeled within the Thy1-YFP-H line belong to the type A subtype of layer 5 pyramidal neurons. **A**. Method of subtype classification. The magnitude of the depolarizing sag in response to hyperpolarizing current injection (difference between 1 and 2) was added to the magnitude of the depolarizing afterpotential which arises upon cessation of hyperpolarizing current (3). All neurons within the present study manifested a positive value of this sum, and were consequently assigned to the type A subtype.

## References

Atapour N & Rosa MGP. (2017). Age-related plasticity of the axon initial segment of cortical pyramidal cells in marmoset monkeys. Neurobiol Aging 57, 95–103.

Baalman KL, Cotton RJ, Rasband SN & Rasband MN. (2013). Blast wave exposure impairs memory and decreases axon initial segment length. J Neurotrauma 30, 741–751.

Basso E, Fante L, Fowlkes J, Petronilli V, Forte MA & Bernardi P. (2005). Properties of the permeability transition pore in mitochondria devoid of Cyclophilin D. J Biol Chem 280, 18558–18561.

Blennow K, Hardy J & Zetterberg H. (2012). The neuropathology and neurobiology of traumatic brain injury. Neuron 76, 886–899.

Brocard C, Plantier V, Boulenguez P, Liabeuf S, Bouhadfane M, Viallat-Lieutaud A, Vinay L & Brocard F. (2016). Cleavage of Na(+) channels by calpain increases persistent Na(+) current and promotes spasticity after spinal cord injury. Nat Med 22, 404–411.

Dembrow NC, Chitwood RA & Johnston D. (2010). Projection-specific neuromodulation of medial prefrontal cortex neurons. J Neurosci 30, 16922–16937.

Evans MD, Sammons RP, Lebron S, Dumitrescu AS, Watkins TB, Uebele VN, Renger JJ & Grubb MS. (2013). Calcineurin signaling mediates activity-dependent relocation of the axon initial segment. J Neurosci 33, 6950–6963.

Favero M, Sotuyo NP, Lopez E, Kearney JA & Goldberg EM. (2018). A Transient Developmental Window of Fast-Spiking Interneuron Dysfunction in a Mouse Model of Dravet Syndrome. J Neurosci 38, 7912–7927.

Feng G, Mellor RH, Bernstein M, Keller-Peck C, Nguyen QT, Wallace M, Nerbonne JM, Lichtman JW & Sanes JR. (2000). Imaging neuronal subsets in transgenic mice expressing multiple spectral variants of GFP. Neuron 28, 41–51.

Forte M, Gold BG, Marracci G, Chaudhary P, Basso E, Johnsen D, Yu X, Fowlkes J, Rahder M, Stem K, Bernardi P & Bourdette D. (2007). Cyclophilin D inactivation protects axons in experimental autoimmune encephalomyelitis, an animal model of multiple sclerosis. Proceedings of the National Academy of Sciences of the United States of America 104, 7558–7563.

Gee S, Ellwood I, Patel T, Luongo F, Deisseroth K & Sohal VS. (2012). Synaptic activity unmasks dopamine D2 receptor modulation of a specific class of layer V pyramidal neurons in prefrontal cortex. J Neurosci 32, 4959–4971.

Goethals S & Brette R. (2020). Theoretical relation between axon initial segment geometry and excitability. Elife 9.

Gomis-Rüth S, Wierenga CJ & Bradke F. (2008). Plasticity of polarization: changing dendrites into axons in neurons integrated in neuronal circuits. Curr Biol 18, 992–1000.

Greer JE, Hånell A, McGinn MJ & Povlishock JT. (2013). Mild traumatic brain injury in the mouse induces axotomy primarily within the axon initial segment. Acta Neuropathol 126, 59–74.

Greer JE, Povlishock JT & Jacobs KM. (2012). Electrophysiological abnormalities in both axotomized and nonaxotomized pyramidal neurons following mild traumatic brain injury. J Neurosci 32, 6682–6687.

Grubb MS & Burrone J. (2010). Activity-dependent relocation of the axon initial segment fine-tunes neuronal excitability. Nature 465, 1070–1074.

Gulledge AT & Bravo JJ. (2016). Neuron Morphology Influences Axon Initial Segment Plasticity. eNeuro 3.

Gutzmann A, Ergül N, Grossmann R, Schultz C, Wahle P & Engelhardt M. (2014). A period of structural plasticity at the axon initial segment in developing visual cortex. Front Neuroanat 8, 11.

Hamada MS, Goethals S, de Vries SI, Brette R & Kole MH. (2016). Covariation of axon initial segment location and dendritic tree normalizes the somatic action potential. Proc Natl Acad Sci U S A 113, 14841–14846.

Hånell A, Greer JE & Jacobs KM. (2015). Increased Network Excitability Due to Altered Synaptic Inputs to Neocortical Layer V Intact and Axotomized Pyramidal Neurons after Mild Traumatic Brain Injury. J Neurotrauma 32, 1590–1598.

Hedstrom KL, Ogawa Y & Rasband MN. (2008). AnkyrinG is required for maintenance of the axon initial segment and neuronal polarity. J Cell Biol 183, 635–640.

Hilscher MM, Leão RN, Edwards SJ, Leão KE & Kullander K. (2017). Chrna2-Martinotti Cells Synchronize Layer 5 Type A Pyramidal Cells via Rebound Excitation. PLoS Biol 15, e2001392.

Hinman JD, Rasband MN & Carmichael ST. (2013). Remodeling of the axon initial segment after focal cortical and white matter stroke. Stroke 44, 182–189.

Hu W, Tian C, Li T, Yang M, Hou H & Shu Y. (2009). Distinct contributions of Na(v)1.6 and Na(v)1.2 in action potential initiation and backpropagation. Nat Neurosci 12, 996–1002.

Iljazi A, Ashina H, Al-Khazali HM, Lipton RB, Ashina M, Schytz HW & Ashina S. (2020). Post-Traumatic Stress Disorder After Traumatic Brain Injury-A Systematic Review and Meta-Analysis. Neurol Sci 41, 2737–2746.

Jamann N, Dannehl D, Lehmann N, Wagener R, Thielemann C, Schultz C, Staiger J, Kole MHP & Engelhardt M. (2021). Sensory input drives rapid homeostatic scaling of the axon initial segment in mouse barrel cortex. Nat Commun 12, 23.

Khaliq ZM & Raman IM. (2006). Relative contributions of axonal and somatic Na channels to action potential initiation in cerebellar Purkinje neurons. J Neurosci 26, 1935–1944.

Kole MH, Ilschner SU, Kampa BM, Williams SR, Ruben PC & Stuart GJ. (2008). Action potential generation requires a high sodium channel density in the axon initial segment. Nat Neurosci 11, 178–186.

Kole MH & Stuart GJ. (2012). Signal processing in the axon initial segment. Neuron 73, 235–247.

Kress GJ, Dowling MJ, Meeks JP & Mennerick S. (2008). High threshold, proximal initiation, and slow conduction velocity of action potentials in dentate granule neuron mossy fibers. J Neurophysiol 100, 281–291.

Kuba H, Oichi Y & Ohmori H. (2010). Presynaptic activity regulates Na(+) channel distribution at the axon initial segment. Nature 465, 1075–1078.

Kuba H, Yamada R, Ishiguro G & Adachi R. (2015). Redistribution of Kv1 and Kv7 enhances neuronal excitability during structural axon initial segment plasticity. Nat Commun 6, 8815.

Lazarov E, Dannemeyer M, Feulner B, Enderlein J, Gutnick MJ, Wolf F & Neef A. (2018). An axon initial segment is required for temporal precision in action potential encoding by neuronal populations. Science advances 4, eaau8621.

Lee AT, Gee SM, Vogt D, Patel T, Rubenstein JL & Sohal VS. (2014). Pyramidal neurons in prefrontal cortex receive subtype-specific forms of excitation and inhibition. Neuron 81, 61–68.

Marin MA, Ziburkus J, Jankowsky J & Rasband MN. (2016). Amyloid-β plaques disrupt axon initial segments. Exp Neurol 281, 93–98.

McGinn MJ, Kelley BJ, Akinyi L, Oli MW, Liu MC, Hayes RL, Wang KK & Povlishock JT. (2009). Biochemical, structural, and biomarker evidence for calpain-mediated cytoskeletal change after diffuse brain injury uncomplicated by contusion. J Neuropathol Exp Neurol 68, 241–249.

Meeks JP & Mennerick S. (2007). Action potential initiation and propagation in CA3 pyramidal axons. J Neurophysiol 97, 3460–3472.

Palmer LM & Stuart GJ. (2006). Site of action potential initiation in layer 5 pyramidal neurons. J Neurosci 26, 1854–1863.

Porrero C, Rubio-Garrido P, Avendaño C & Clascá F. (2010). Mapping of fluorescent protein-expressing neurons and axon pathways in adult and developing Thy1-eYFP-H transgenic mice. Brain Res 1345, 59–72.

Povlishock JT & Christman CW. (1995). The pathobiology of traumatically induced axonal injury in animals and humans: a review of current thoughts. J Neurotrauma 12, 555–564.

Prins M, Greco T, Alexander D & Giza CC. (2013). The pathophysiology of traumatic brain injury at a glance. Dis Model Mech 6, 1307–1315.

Rowe RK, Griffiths DR & Lifshitz J. (2016). Midline (Central) Fluid Percussion Model of Traumatic Brain Injury. Methods Mol Biol 1462, 211–230.

Schafer DP, Jha S, Liu F, Akella T, McCullough LD & Rasband MN. (2009). Disruption of the axon initial segment cytoskeleton is a new mechanism for neuronal injury. J Neurosci 29, 13242–13254.

Spiegel DP, Reynaud A, Ruiz T, Laguë-Beauvais M, Hess R & Farivar R. (2016). First- and second-order contrast sensitivity functions reveal disrupted visual processing following mild traumatic brain injury. Vision Res 122, 43–50.

Stuart G, Spruston N, Sakmann B & Häusser M. (1997). Action potential initiation and backpropagation in neurons of the mammalian CNS. Trends Neurosci 20, 125–131.

Sun J & Jacobs KM. (2016). Knockout of Cyclophilin-D Provides Partial Amelioration of Intrinsic and Synaptic Properties Altered by Mild Traumatic Brain Injury. Front Syst Neurosci 10, 63.

Sun J & Jacobs KM. (2017). Neurophysiological Study of Traumatic Brain Injury: To Slice or Not to Slice. Open Access Journal of Neurology & Neurosurgery 2, 555600.

Vascak M, Sun J, Baer M, Jacobs KM & Povlishock JT. (2017). Mild Traumatic Brain Injury Evokes Pyramidal Neuron Axon Initial Segment Plasticity and Diffuse Presynaptic Inhibitory Terminal Loss. Front Cell Neurosci 11, 157.

von Reyn CR, Spaethling JM, Mesfin MN, Ma M, Neumar RW, Smith DH, Siman R & Meaney DF. (2009). Calpain mediates proteolysis of the voltage-gated sodium channel alpha-subunit. J Neurosci 29, 10350–10356.

Wefelmeyer W, Cattaert D & Burrone J. (2015). Activity-dependent mismatch between axo-axonic synapses and the axon initial segment controls neuronal output. Proc Natl Acad Sci U S A 112, 9757–9762.

Yermakov LM, Drouet DE, Griggs RB, Elased KM & Susuki K. (2018). Type 2 Diabetes Leads to Axon Initial Segment Shortening in db/db Mice. Front Cell Neurosci 12, 146.

Zhao Y, Wu X, Chen X, Li J, Tian C, Chen J, Xiao C, Zhong G & He S. (2020). Calcineurin Signaling Mediates Disruption of the Axon Initial Segment Cytoskeleton after Injury. iScience 23, 100880.

