## Supplemental Figure 1 for "Concussive head trauma perturbs axon initial segment function in axotomized and intact layer 5 pyramidal neurons"

**Supplemental Fig. 1**

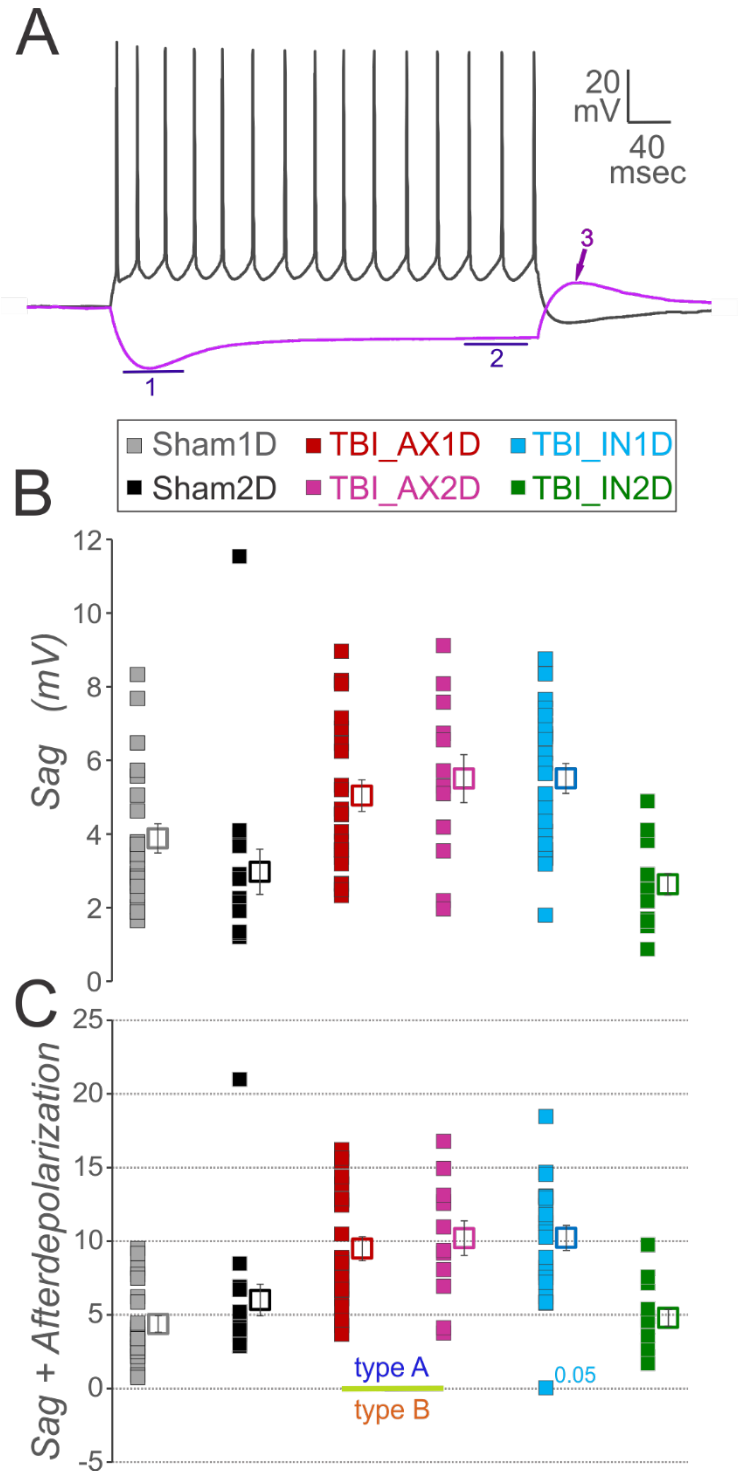


Supplemental figure 1. $I_h$ current and Type A vs Type B neurons. The $I_h$ current was recognized as a depolarizing sag in response to hyperpolarizing current injection. **A.** Examples showing how $I_h$ and the mean of the steady state afterdepolarization (~ 50 msec after the end of current injection pulse) were measured. Sag was measured as the difference between the steady-state hyperpolarization at end of pulse (marked as 2) and the peak sag (marked as 1). The afterdepolarization, to which calcium T-currents contribute, was measured at the peak within

70 msec of the pulse ending minus the resting level prior to the pulse (arrow marked as 3). **B.** Sag measured for individual neurons in each WT subject group (filled symbols), and the mean for the group (open symbols) with SEM in error bars. A 1-way ANOVA showed a significant effect of subject group and Bonferroni post-hoc showed that groups AX1D, AX2D, and IN1D had a significantly larger sag compared to IN2D and sham2D. **C.** Previous studies have shown that adding the afterdepolarization to the sag results in a measure that distinguishes Type A versus Type B neurons. These two distinct layer 5 pyramidal neuron groups have previously been shown to differ in: their projection sites, their dendritic and axonal morphologies, and a number of electrophysiological characteristics beyond just $I_h$ and T-channel currents (Gee *et al.*, 2012). While most previous studies on type A vs type B were conducted in prefrontal cortex, we have found that within somatosensory cortex of mice that the presence of $I_h$ begins when this summed measure is above 0 (data not shown). All of the YFP+ pyramidal neurons in this study had some $I_h$ and were above zero for the sum of sag + afterdepolariztion. Thus, they all fall into the type A category. Similar to the measurement of $I_h$ and likely due to that measure, there was also a significant difference among groups on the sum of sag and the afterdepolarization, based on a 1-way ANOVA. Again, groups AX1D, AX2D, and IN1D were significantly larger compared to IN2D and sham2D. N = 22 Sham1D, 16 Sham2D, 23 AX1D, 12 AX2D, 22 IN1D, and 14 IN2D neurons.
